# Full genome characterization of Laem Singh virus (LSNV) in shrimp *Penaeus monodon*

**DOI:** 10.1101/2020.07.25.221432

**Authors:** Suparat Taengchaiyaphum, Jiraporn Srisala, Piyachat Sanguanrut, Chalermporn Ongvarrasopone, Timothy W. Flegel, Kallaya Sritunyalucksana

## Abstract

Laem Singh virus (LSNV) was discovered in 2006 and proposed as a necessary but insufficient cause of retarded growth in the giant tiger shrimp *Penaeus monodon.* Its closest relatives were plant viruses including an unassigned *Sobemovirus* and viruses in the family *Luteoviridae*. During succeeding years, attempts to obtain the full LSNV genome sequence by genome walking failed. However, recent publication of the full sequence of Wenzhou shrimp virus 9 (WZSV 9) at GenBank revealed that LSNV sequences in our database shared 99% sequence identity with it. Thus, we hypothesized that LSNV and WZSV 9 were different isolates of the same virus species. Here we confirm that hypothesis by cloning and sequencing of the full genome of LSNV from *P. monodon* and by showing that it consists of two fragments each with 99% identity to the matching fragments of WZSV.

## INTRODUCTION

Laem Singh virus (LSNV) was discovered in 2006 (Sritunyalucksana, et al., 2006). This was accomplished by using differential centrifugation followed by density-gradient centrifugation, to obtain potential viral bands that were then subjected to total nucleic acid extraction (i. e., RNA plus DNA). The extracts were, in turn, used as templates for a random-primer, reverse-transcriptase, polymerase chain reaction (RT-PCR) process to produce cDNA from any RNA present, followed by random primer PCR of the resulting mixture containing both cDNA and DNA. The resulting cDNA-DNA mix was subjected to shotgun cloning. Resulting amplicon sequences were compared with the GenBank database to identify potentially novel, viral sequences. Candidate sequences were used to design specific primers for use in PCR and RT-PCR testing of the original gradient-centrifugation extracts and of normal and growth-retarded shrimp. Ultimately, LSNV was proposed as a necessary but insufficient cause of retarded growth (called monodon slow growth syndrome or MSGS) in the giant tiger shrimp *Penaeus monodon* (Sritunyalucksana, et al., 2006). The missing component cause of MSGS has been reported to be associated with retinopathy (Pratoomthai, et al., 2008) and an integrase-containing element (Panphut, et al., 2011). MSGS was a major reason for the switch from indigenous *P. monodon* to exotic *P. vannamei* (unaffected by LSNV infection) as the major cultivated species in Thailand from around 2002 onwards. LSNV has been reported from *P. monodon* in other countries (Prakasha, et al., 2007; Sittidilokratna, et al., 2009) but not as associated with retarded growth.

A sufficient portion of the LSNV genome was obtained to identify its closest relatives to be plant viruses including an unassigned *Sobemovirus* and viruses in the family *Lutoviridae*. During succeeding years, we failed in our attempts to obtain the full LSNV genome sequence by genome walking. However, after deposit of the full sequence of Wenzhou shrimp virus 9 (WZSV 9) at GenBank (Shi, et al., 2016), we discovered that the LSNV sequences in our database shared 99% sequence identity with homologous regions in the genome of WZSV 9. We also realized that our genome walking work may have failed because the WZSV 9 genome was in two fragments, unlike the viruses in the genus *Sobemovirus* and the family *Luteoviridae*. Thus, we hypothesized that LSNV and WZSV 9 were different isolates of the same virus. To test this hypothesis, we designed primers based on the two WZSV 9 fragments in order to amplify and sequence the full genome of LSNV from infected Thai shrimp for comparison to that of WZSV 9. Results herein confirm that hypothesis by showing that the full genome of LSNV is in two fragments, each of which have 99% sequence identity to the published sequence of WZSV 9.

## MATERIALS AND METHODS

### 1. Shrimp samples

*Penaeus monodon* broodstock specimens of approximately 80-100 g body weight were obtained from Shrimp Genetic Improvement Center (SGIC, BIOTEC), Surat Thani province, Thailand.

### 2. LSNV viral particle purification

For LSNV purification, 10 g whole gills from LSNV-infected shrimp were homogenized in 20 ml iced-cooled TN buffer (20 mM Tris-HCl (pH 7.5), 400 mM NaCl). The suspension was clarified twice by low-speed centrifugation. The supernatant was transferred to a new tube and centrifuged at 200,000 × g for 2 h at 4°C using a ultracentrifuge. After centrifugation, the pellet was collected and resuspended in 400 μl TN buffer before the suspension was overlayered onto a 15– 60% continuous sucrose gradient and centrifuged at 286,000 × g, 4°C for 2 h. The band at approximately 45-60% sucrose was collected and washed with 5 ml of TN buffer before centrifugation at 286,000 × g, 4°C for 2 h. Finally, the pellet was dissolved in small volume of TN buffer and kept at 4°C. The LSNV particle size was checked by transmission electron microscopy (TEM) using the negative staining technique, i.e., virus particles stained with 2% phosphotungstic acid (PTA), pH 7.0 on copper, formvar-coated grids.

### 3. Total RNA extraction

Lymphoid organs (a prime target of LSNV) (Pratoomthai, et al., 2008; Sritunyalucksana, et al., 2006) were removed by dissection and homogenized in Trizol reagent (Invitrogen) prior to total RNA extraction using the chloroform-isopropyl alcohol method. The total RNA was collected by centrifugation and washed with 75% ethanol. The RNA pellet was resuspended in DNase-RNase free water (Gibco) and then further digested DNase I (NEB) following the manufacturer’s protocol. Subsequently, the sample was re-extracted by the same method. Total RNA concentration was determined by Nanodrop spectrophotometer (Thermo Scientific). The RNA stock was kept at −80 °C until used.

### 4. RACE amplification

The first PCR reaction was performed in 25 μl containing 0. 4 μM of A1-991 primer and S1-991 primer, 0. 2 mM of dNTP, 1X BD Advantage™ 2 PCR buffer, 1X BD Advantage™ 2 polymerase Mix, using 3 μl of diluted (1:10) cDNA as the template. The PCR reaction protocol consisted of 94°C for 5 min followed by 25 cycles of 94°C for 30 sec, 55°C for 30 sec, 72°C for 2 min and final extension at 72°C for 7 min. The PCR products were analyzed by 1. 2% agarose gel electrophoresis and UV light visualization. If necessary, nested PCR reactions were carried out in 25 μl containing 0.4 μM of A2-991 primer and S2-991 primer, 0.2 mM of dNTP, 1X BD Advantage™2 PCR buffer, 1X BD Advantage™2 polymerase Mix, using 5 μl of diluted (1:10) PCR product from the first PCR reaction. The PCR protocol consisted of 94°C for 5 min followed by 20 cycles of 94°C for 30 sec, 55°C for 30 sec, 72°C for 3 min and final extension at 72°C for 7 min. The final PCR products were analyzed by 1. 2% agarose gel electrophoresis and UV light visualization. All primers are shown in **Table 1**.

**Table 1.**
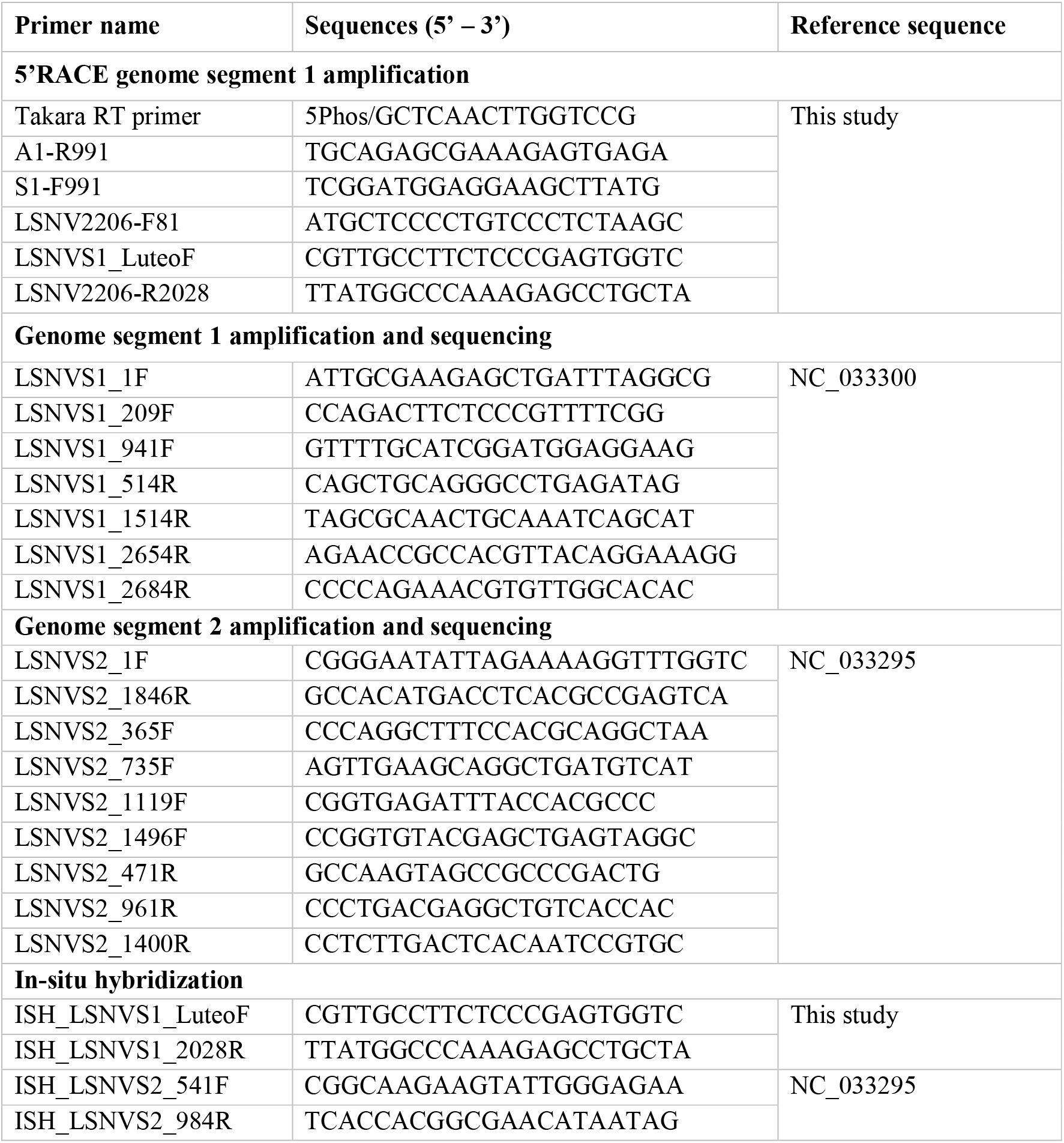
List of primers used in this study.

DNA amplicons were processed by gel purification. Briefly DNA bands were sliced from gels and purified using a QIAquick Gel purification kit (QIAGEN, USA) following the instruction manual. Cloning employed pGEM-T-Easy vector and was carried out in 10 μl containing 1X rapid ligation buffer [30 mM Tris-HCl (pH 7.8), 10 mM MgCl2,10 mM DTT, 1 mM ATP and 5% polyethylene glycol], 2 μl of gel-purified PCR product, 50 ng of pGEM-T-Easy vector and 3 U of T4 DNA ligase. The mixture was incubated at 4°C overnight. The ligated products were transformed into *E. coli* JM109.

### 5. RT-PCR

To characterize LSNV RNA segments, one-step RT-PCR was carried out using a One-step RT-PCR detection kit (Invitrogen). Briefly, 1X PCR master mix containing 0.4 μM forward and reverse primers (primers are shown in **Table 1**), 1U Platinum™*Taq* polymerase, and 100 ng of total RNA was subjected to RT-PCR using a PCR thermocycler (Bio-Rad). The RT-PCR protocol consisted of 50°C or 30 min followed by initial denaturation at 94°C for 2 min, then 30 cycles of PCR including 94°C for 15 s, 58°C for 30 s, 72°C for 2 min and final extension at 72°C for 5 min. The PCR amplicons were separated by 1.5% agarose gel electrophoresis, visualized by UV illuminator and recorded by EZ gel documentation (Bio-Rad).

For PCR amplification and DNA sequencing, total mRNA was subjected to cDNA synthesis using SuperScript™ III One-Step RT-PCR System with Platinum™ *Taq* DNA Polymerase (Invitrogen). The PCR products were separated by 1. 5% agarose gel electrophoresis. Subsequently, each PCR product was excised from the agarose gel and then subjected to further PCR product purification using a Gel/PCR purification kit (GeneAid). The concentration of purified PCR-amplicon DNA was determined by Nanodrop spectrophotometer prior to DNA sequencing analysis by Macrogen DNA sequencing service (Macrogen, Korea). Three individual PCR products from 3 individual shrimps were sent for DNA sequencing.

### 6. DNA sequencing and phylogenetic analysis

The clones were sequenced by Macrogen Inc. (Korea). Overlapping sequences were compiled using CAP3 sequence Assembly Program (available online at http://pbil.univ-lyon1.fr/cap3.php). BCM search launcher program was used to identify open reading frames of nucleotide sequences (available online at http://searchlauncher.bcm.tmc.edu/seq-util/Options/sixframe.html). Nucleotide and deduced amino acid sequences were compared to other known sequences in the GenBank database using BlastX (available online at http://www.ncbi.nlm.nih.gov/BLAST/). The deduced amino acid sequences were aligned with other proteins belonging to the viral RdRp group using ClustalW 1. 7 multiple sequence alignment (available online at www.ebi.ac.uk/clustalw/). The phylogenetic tree was constructed using MEGA-X program. Tree topology was evaluated by bootstrap analysis using the neighbor-joining method with default parameters for 1000 replicates.

For PCR product sequencing, gel/ PCR purification was carried out to obtain single PCR fragments after RT-PCR. The purified PCR products were sent for sequencing using several forward and reverse primers to cover all genome regions. Retrieved DNA sequences were trimmed and re-assembled to obtain full-length sequences. Prediction of open reading frames and deduced amino acid sequence alignments were performed using ORF finder (https://www.ncbi.nlm.nih.gov/orffinder/) and BlastP analysis (http://www.ncbi.nlm.nih.gov/BLAST/).

### 7. In-situ hybridization (ISH)

Two *P. monodon* broodstock were processed for histological analysis according to standard methods (Bell Lightner, 1988). Briefly tissues were fixed in Davidson’ s fixative solution for 24 h before transfer to 70% ethanol followed by processing of the cephalothorax to prepare adjacent paraffin tissue sections for hematoxylin and eosin (H&E) staining and for ISH. For ISH, de-paraffinized tissue sections were incubated with 5 μg/ ml of Proteinase K in 1xTNE buffer at 37°C for 15 min pre-hybridization solution. Subsequently, tissue was immersed in 0.5M EDTA for 1 h before transfer to 0.4% formaldehyde solution for 5 min prior to incubated with pre-hybridization buffer at 37°C for 30 min. This was followed by application of the DNA probe solution in hybridization buffer for overnight incubation at 42°C in a moisture chamber. Negative controls consisted of adjacent sections treated with hybridization buffer containing no probe. Next, tissue sections were washed several times with washing buffers before incubation with NBT/BCIP substrate (Roche) from 30 min to overnight at 4°C. Subsequently, tissue was counter-stained with 0.5% Bismarck brown dye for 15-30 min prior to washing with tap water, dehydration and mounting with permount. The positive ISH signals were assessed by light microscopy using 10X to 40x objective lenses.

Probes were prepared by PCR amplification using a PCR-DIG labelling kit (Roche) as directed by the manufacturer. Lists of primers used for probe amplification are shown in **Table 1**. PCR products were purified using a gel/PCR extraction kit and DNA concentration was determined by Nanodrop spectrophotometer. Specific probes (150 ng) were used to detect each of the two different viral genome fragments.

## RESULTS

### 1. LSNV-like particles purified by sucrose gradient centrifugation

Sucrose gradient centrifugation of tissues from shrimp positive for LSNV by RT-PCR resulted in two distinct layers between 54 and 58% sucrose concentration (**Figure 1A**). TEM revealed the presence of viral-like particles ranging from 23-33 nm relative to accompanying Nano-bead particles (**Figure 1B**). RT-PCR detection of LSNV-related sequences using total RNA templates extracted separately from the two bands gave positive amplicons from both bands (**Figure 1C**, L1 and L2) indicating that both bands contained LSNV RNA. In addition, RNase A digestion of the RNA templates prior to RT-PCR testing resulted in no amplicons. This confirmed that the amplicons resulted from the presence of LSNV RNA.

**Figure 1.**
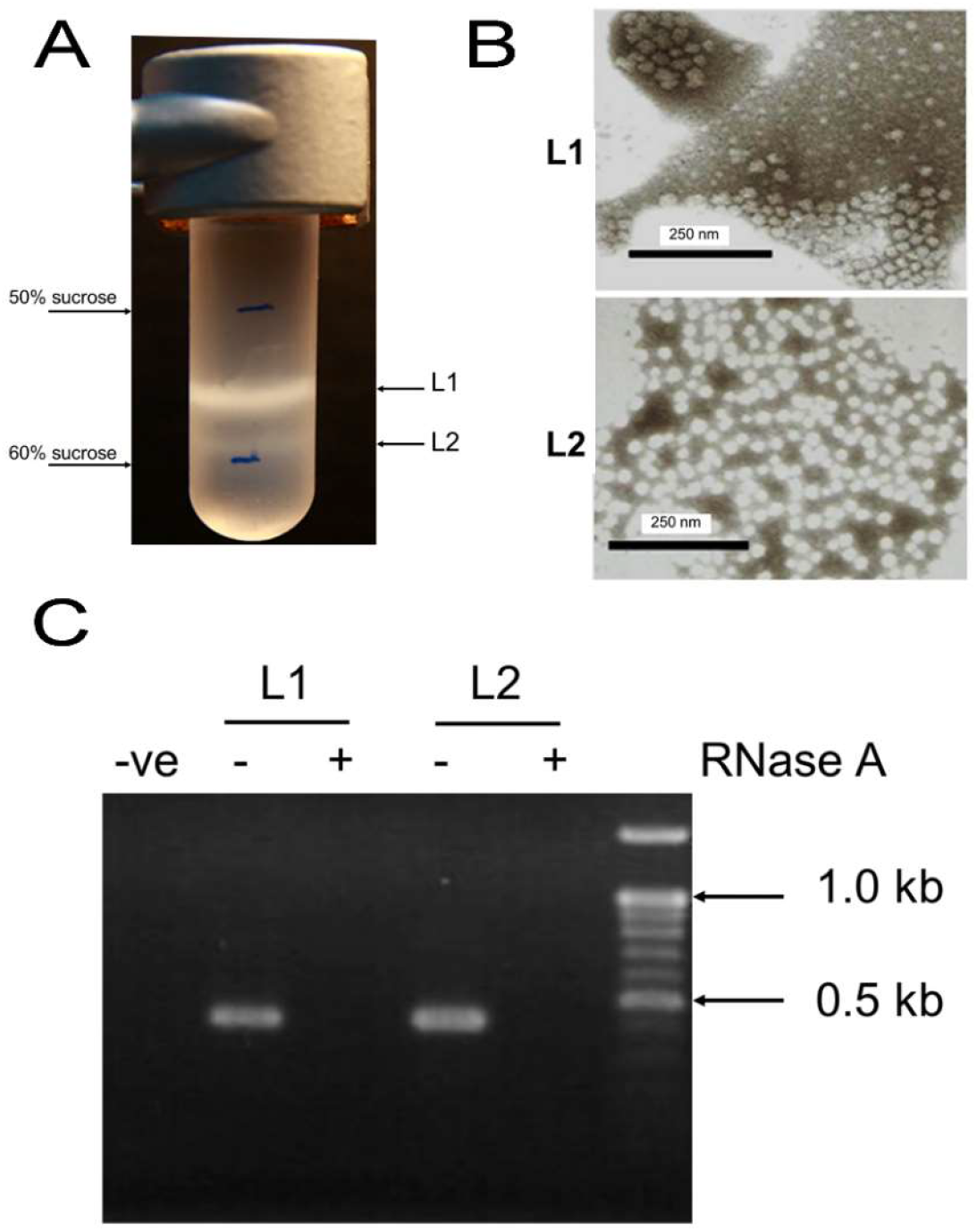
Purification of LSNV from infected *P. monodon* gill tissue. (**A**) Sucrose density gradient centrifugation tube showing 2 layers between 54 and 58%. (**B**) TEM showing icosahedral viral-like particles with diameters ranging from 23-33 nm. (**C**) Agarose gel showing RT-PCR amplicons using templates with and without RNase treatment for both

### 2. LSNV genome segments and sequence analysis

RACE amplifications and RT-PCR detection of full-length LSNV genome fragments were carried out. Two fragments were obtained. One was 2,206 bp in length, and sequence analysis using BlastN showed 99% similarity to Wenzhou virus 9 (WZSV 9) genome fragment 1 (GenBank accession no. NC_033300). RT-PCR detection based on specific primers designed from RACE and WZSV 9 fragment 1 gave a positive single amplicon in infected shrimp (**Figure 2A, left**). Multiple sequence alignment also indicated 99% identity between WZSV 9 genome fragment 1 and the LSNV-RACE amplicon and the RT-PCR amplicon by DNA sequencing **(supplementary Figure S1**).

**Figure 2.**
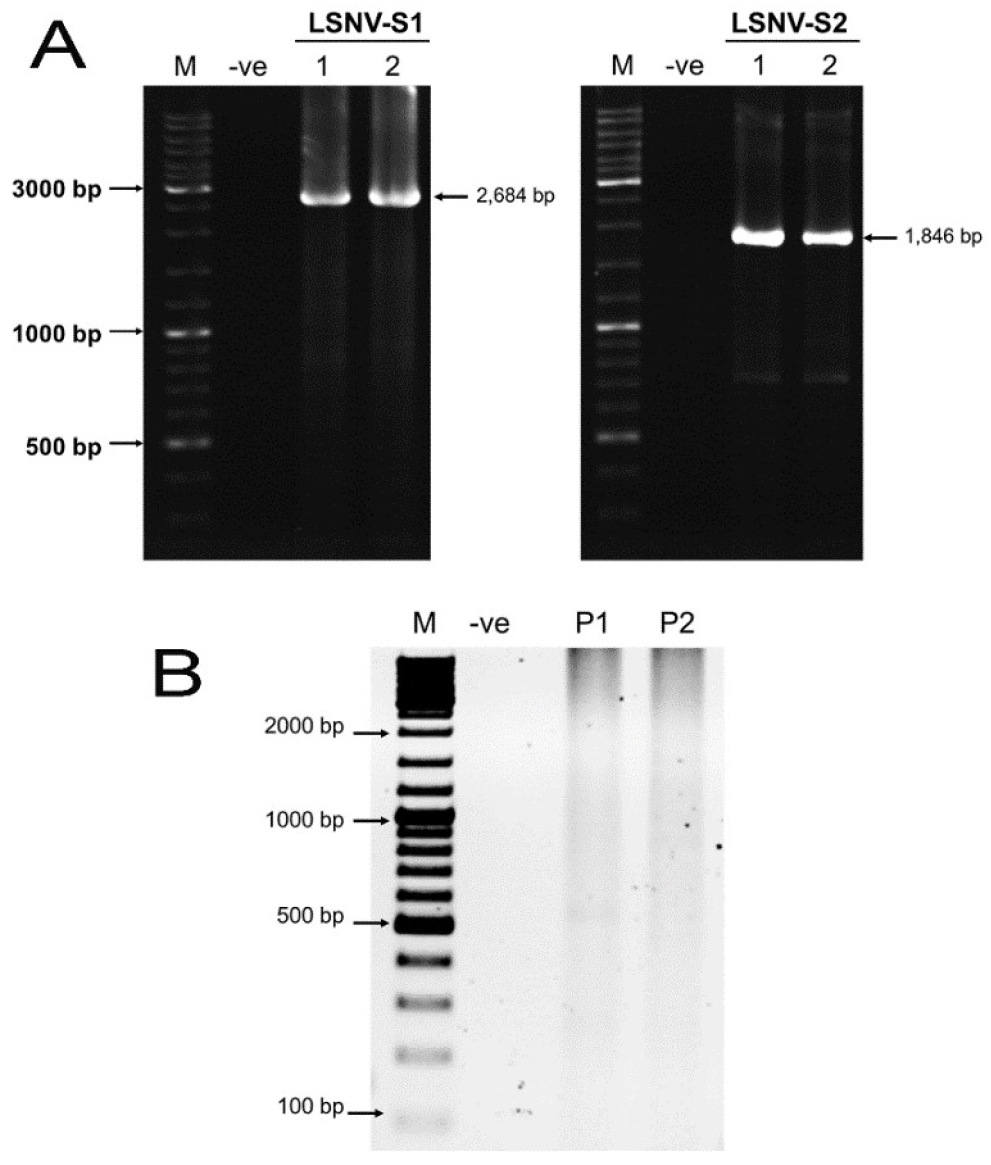
RT-PCR detection of LSNV genome segments. (A) Agarose gels showing full-length amplification of LSNV genome segments of 2,684 bp for segment 1 and 1,846 bp using specific primers using template RNA from LSNV-infected shrimp. (B) RT-PCR testing by using overlapping primers between two LSNV genome segments. Absence of amplicons was observed when using different pairs of primer flanking between genome segment 1 and segment 2 (primer set 1 (P1); LSNVS1_Luteo-F + LSNVS2-1810R, and primer set 2 (P2); LSNVS1_941F + LSNVS2-471R).

The second RT-PCR amplification using the same samples but with specific primers designed from WZSV 9 genome fragment 2 (performed in parallel) gave a positive amplicon of 1846 bp (**Figure 2A, Right**) with 99% sequence identity to that of WZSV 9 genome fragment 2 **(supplement figure S2)**. No amplicons arose when reactions were carried out using any forward primer from genome fragment 1 with any reverse primer from genome segment 2 (**Figure 2B**), confirming that the two fragments in LSNV were not contiguous, i.e., that the genome was bi-partite.

Prediction of open reading frames in genome fragments 1 and 2 was predicted using online analysis tools. Two possible open reading frames were found. In PCR amplicon 1 (called LSNV-F1_PCR), this included one putative protease-like protein of 453 amino acids and one putative RNA-dependent RNA polymerase (RdRp) of 382 amino acids (**Figure 3**). ORF prediction for PCR amplicon 2 (called LSNV-F2_PCR) revealed only one protein, a putative viral capsid protein of 585 amino acids (**Figure 3B**). Multiple sequence alignment of deduced amino acids from both fragments revealed 99% amino acid identity with the matching ORF of the 2-fragment WZSV 9 genome **(supplement figure S3-S5**). Most of the top hits in the phylogenetic analysis were from the plant, *Sobemovirus*-like group. Overall, the results indicated that LSNV and WZSV 9 were isolates of the same virus species, with a genome organization similar to that of plant *Sobemovirus* (i. e., 3 major proteins genes including a hypothetical protein, an RdRp and a capsid protein). The only major difference was that the *Sobemovirus* genome consists of single ssRNA fragment while the genomes of LSNV and WZSV 9 are divided into 2 fragments (i.e., bi-partite) (**Figure 3C**).

**Figure 3.**
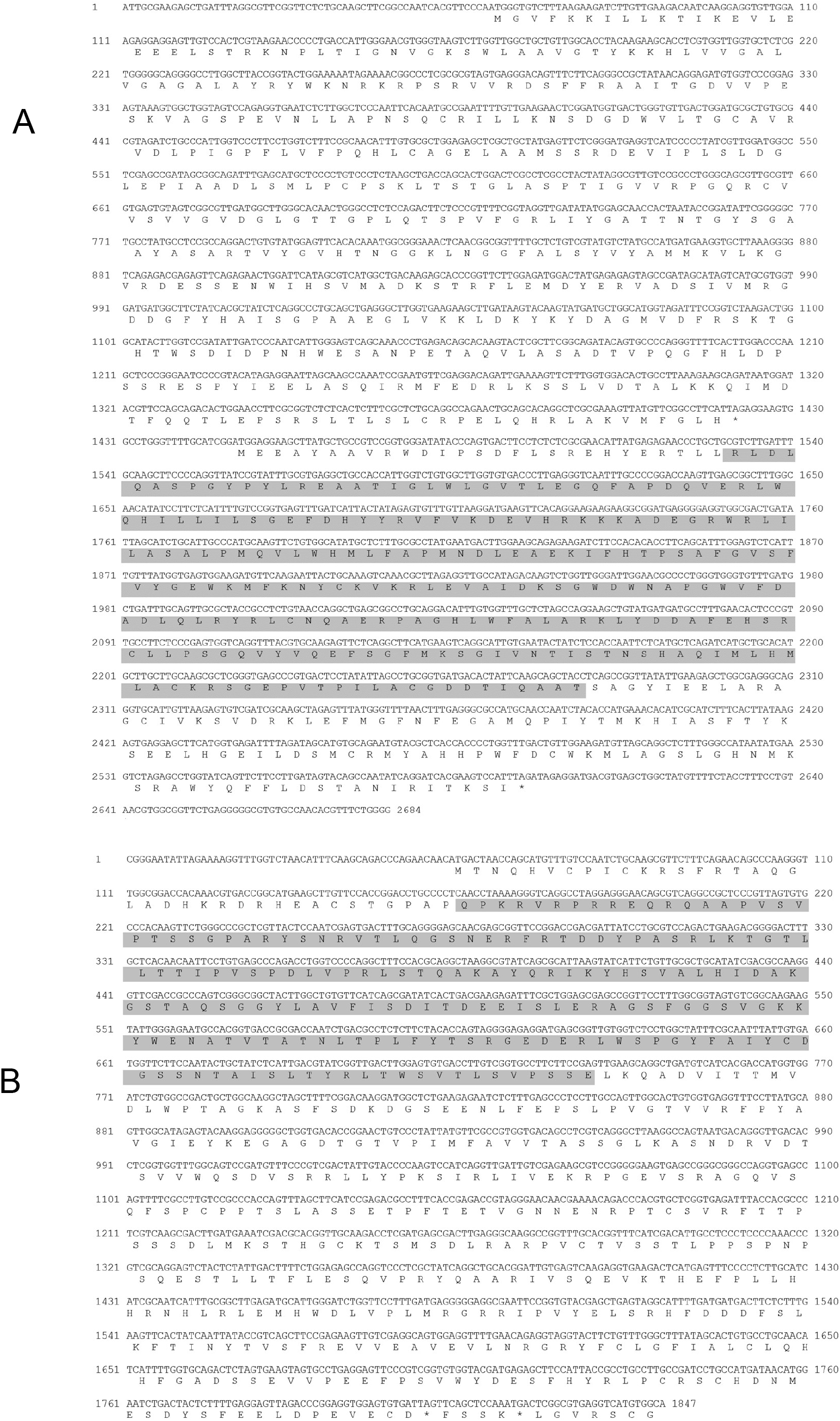
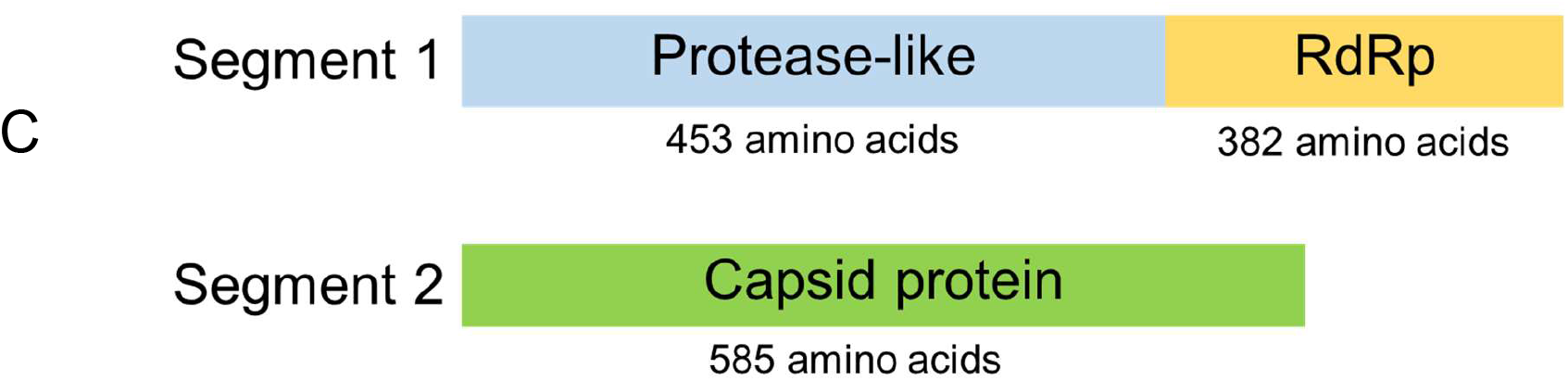
cDNA sequences of LSNV RT-PCR amplicons and predicted open reading frames. (**A**) Sequence of LSNV genome segment 1 with two predicted-ORF of 453 and 382 deduced amino acids related to a protease-like protein and an RdRP respectively, based on BlastP analysis. The shaded box indicates a conserved region corresponding to the reverse transcriptase-RNA-dependent RNA polymerase-like superfamily. (**B**) DNA sequence of LSNV genome segment 2 with a single predicted-ORF of 585 deduced amino acids and showing sequence similarity to capsid proteins. The shaded box indicates a conserved region corresponding to viral capsid protein family. (**C**) Schematic diagram showing the structure of the LSNV genome segments and predicted proteins compositions (figure not to scale).

### 3. Phylogenetic analysis

Blast P analysis using amino acid sequences of predicted proteins from the two LSNV genome segments detected from *P. monodon* was carried out to determine their relationships to other proteins in the GenBank database. Two deduced proteins were predicted from genome segment 1 and one from genome segment 2. For each predicted polypeptide, the top 10 hits for most closely related polypeptides obtained from BlastP were collected and aligned using Clustal Omega (Figs. S3-S5). The analysis revealed that all 3 polypeptides predicted from the 3 ORFs in the LSNV genome shared 99-100% identity with the matching polypeptides predicted from the WZSV 9 genome. In contrast, the next most related polypeptides showed only 40% identity or less. We concluded that LSNV and WZSV 9 were representatives of the same viral species.

In the phylogenetic trees for the 3 putative LSNV/WZSV 9 proteins **(Fig. 4A-C)**, the most closely related viruses from a known viral family (40% identity or less) were records for *Sobemovirus*-like viruses in the *Solimoviridae* that currently contains only plant viruses in the genera *Sobemovirus* and *Polemovirus*. Thus, the features of the alignment analysis and the bipartite genome of the LSNV/WZSV 9 species indicated that it could not yet be clearly assigned to any currently defined viral family.

**Figure 4.**
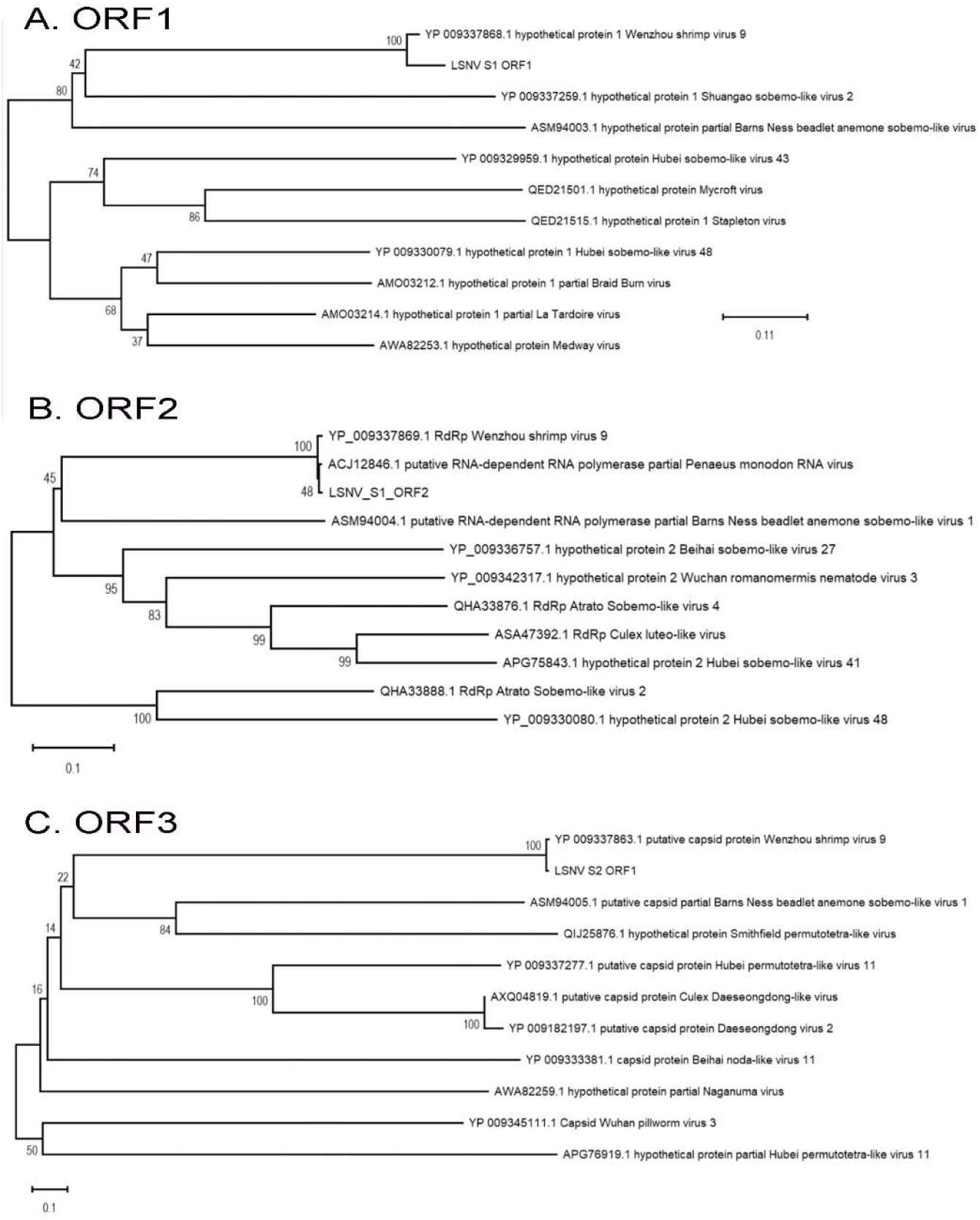
Phylogenetic analysis of predicted polypeptides derived from LSNV genome segment 1 and segment 2. Phylogenetic trees comparing (A) protease-like protein (LSNV_S1, ORF1), (B) RNA-dependent RNA polymerase (LSNV_S1, ORF2), and (C) Capsid protein (LSNV_S2, ORF1) with related viral proteins. The percentage of replicate trees in which the associated taxa clustered together in the bootstrap test (1000 replicates) are shown next to the branches. The tree is drawn to scale, with branch lengths (next to the branches) in the same units as those of the evolutionary distances used to infer the phylogenetic tree. GenBank accession numbers are shown for each viral polypeptide sequence.

### 4. ISH revealed co-localization of LSNV genome fragments in target tissue

Adjacent tissue sections from shrimp positive for LSNV by RT-PCR were subjected to ISH analysis of the lymphoid organ (LO), a known target for LSNV (Sritunyalucksana, et al., 2006). A probe specific for LSNV-F1 was used for one tissue section while a DIG-labeled probe specific for LSNV-F2 was used for the other. The results revealed that the positive ISH reactions for both probes arose from the same location within the LO in the adjacent tissue sections (**Figure 5**). Since the LO is a key target of LSNV in infected shrimp, the positive co-staining of the same tissue regions in infected shrimp, together with the data from RT-PCR and sequencing supported the concept that the LSNV genome consisted of two genome fragments.

**Figure 5.**
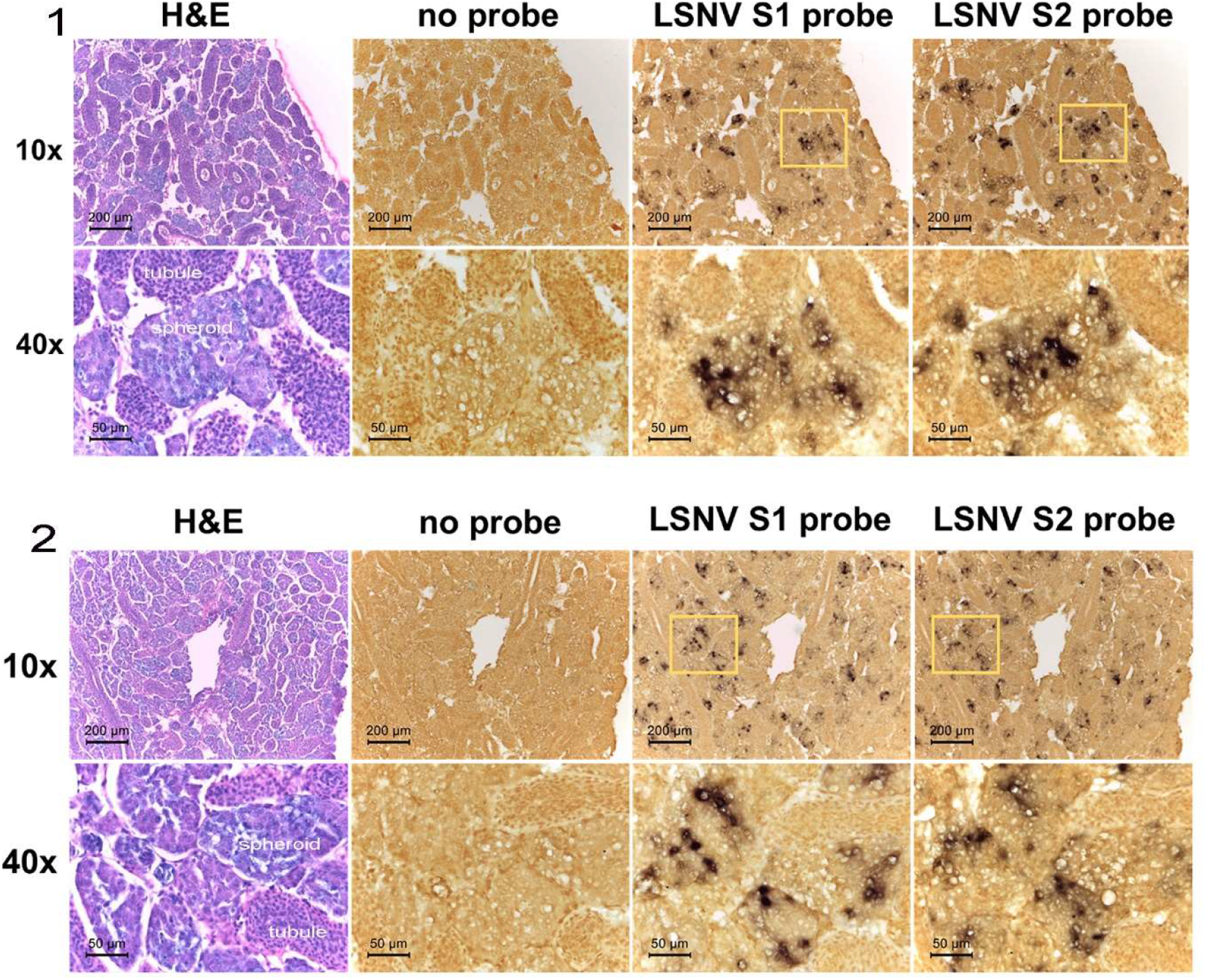
*In-situ* hybridization of LSNV genome segments in infected shrimp *P. Monodon.* Photomicrographs using 10x and 40x objectives of *in-situ* hybridization test results. Photomicrographs in each row were taken from 4 adjacent tissue sections. S1 and S2 probes specific for each LSNV genome segment show dark co-staining regions in spheroids as opposed to tubules of the LO of two individual shrimp specimens (1 & 2) infected with LSNV. The location of zoom-in areas are indicated by yellow boxes.

## Discussion

### LSNV and WZSV 9 are variants of the same viral species

We have shown by genome sequence, genome structure and phylogenetic analysis of the full sequences of LSNV and WZSV 9, that they come from conspecific viral sources. The phylogenetic trees indicated that the most closely related viruses to this species have been reported from insects and other arthropods (Shi, et al., 2016) and been assigned to the *Sobemovirus*-like group (Sõmera, et al., 2015). This concurred with our earlier conclusion that the RdRp sequence of LSNV was most closely related to viruses from the *Luteoviridae* and *Sobemovirus* groups (Sritunyalucksana, et al., 2006).

The family *Sobemoviridae* was initially proposed (Sõmera, et al., 2015) solely based on information from plant pathogens, at least some of which were considered to be transmitted mechanically by insects. Later, the family name was changed to *Solemoviridae* to reflect that it contained the two genera *Sobemovirus* and *Polemovirus* (Sõmera, et al., 2017). However, more recent information suggests that Sobemo-like viruses are not confined to plants.

### *Sobemo-*like viruses widespread in invertebrates

Most of the *Sobemovirus* species were previously believed to be plant pathogens that were mechanically transmitted to plants by insects. However, our review of the literature did not convince us that at least some of these insect vectors might be persistently infected with the Sobemoviruses they carry but show no signs of disease. This is known to occur for vectors of other plant pathogens (Kaur, et al., 2016; Lan, et al., 2016; Rubinstein Czosnek, 1997). At the same time, recent high-throughput sequencing studies with insects and with other invertebrates have revealed that previously unknown *Sobemovirus*-like species may infect many invertebrate hosts (Shi, et al., 2016). These include non-plant-pest insects, spiders, crustaceans, anemones and nematodes. However, the studies are new and based on analysis and assembly of genome fragments. It has not yet been proven for all that they arise from host infectious agents. They also include viruses with both single and bipartite positive-sense RNA genomes.

The fact that the genome of LSNV/WZSV 9 is bipartite contrasts sharply with all viral species currently included in the genus *Sobemovius* in the family *Solemoviridae*. At the very least, this distinction would probably lead to the designation of a new genus, if it were decided that LSNV and other closely related viruses with bi-partite genomes should be included in that family. This would not be without precedence. For example, the family *Tombusviridae* (sister to family *Solemoviridae*) contains two genera, *Umbravirus* with a non-partite genome and *Dianthovirus* with a bi-partite genome (Sõmera, et al., 2015). Alternatively, the bi-partite feature might eventually be used along with other features to erect a separate family for LSNV/WZSV 9.

In arguments for creation of the family *Sobemoviridae* that eventually came to be called *Solemoviridae,* Sõmera stated (Sõmera, et al., 2015) that “ *Polerovirus*, *Enamovirus*, and *Barnavirus* belong to picornavirus-like RdRp superfamily [238]. Hence, there are two big families related to each other—*Luteoviridae* and *Tombusviridae*—but grouped using different main criteria (CP or RdRp homology) as argument for a membership (Figure 5). The genus *Sobemovirus* has phylogenetic relationships with both of them, but it cannot be assigned into one or another family, as it does not share homology with *Luteoviridae* at the CP level or with *Tombusviridae* at the RdRp level. Moreover, genomic sequences of individual sobemovirus species cluster very tightly together. Thus, it is impossible to assign the genus *Sobemovirus* to any existing viral family.”

With LSNV/ WZSV 9, the two most closely-related sequences of the hypothetical protein of (ORF1 and putative protease) are both from the *Solemoviridae* (**Fig. 4A**). One is for Shuangao sobemo-like virus 2 (GenBank YP_009337259)(Shi, et al., 2016)from an insect source and the other, Barns Ness beadlet anemone sobemo-like virus 1 (GenBank ASM94003. 1)(Waldron and Obbard, direct submission). For ORF2 (RdRp) (**Fig. 4B**), the closest record was for Hubai sobemo-like virus 48 (GenBank NC_032233.1) (Shi, et al., 2016) from an insect and the second closest was for Atrato Sobemo-like virus 2 (Nitsche et al., direct submission) also from an insect. For ORF3 (CP) (**Fig. 4C**) in genome fragment 2, the closest record was for Beihai noda-like virus 11 (GenBank YP_009333381.1)(Shi, et al., 2016) from a crustacean and the second closest was for Barns Ness Beadlet anemone sobemo-like virus 1 (ASM94005.1).

For taxonomy, focusing only on the RdRp (ORF2) and CP (ORF3) proteins as described by Sõmera et al. above, the RdRp sequence of LSNV/WZSV 9 matches with the current concept of the family *Solemoviridae*. However, for the CP, the closest matching record was for a pooled crustacean RNA mix while the second record was from an anemone, both from host groups not previously reported to harbor Sobemoviruses. This uncertainty together the explosion in the realm of invertebrate RNA virus discovery (Li, et al., 2015; Shi, et al., 2016) suggest that it will take some time for the taxonomy to settle. This is especially true when considering the distribution of species between the animal and plant kingdoms and whether and how important might be shuttling of the same virus species between infected invertebrate and plant hosts, if it occurs. This is a well-known phenomenon with arboviruses such as Dengue virus, that cause persistent infections without signs of disease in its mosiquito carriers by mechanisms that have only recently been discovered (Tassetto, et al., 2019; Whitfield, et al., 2017). Thus, it seems possible that similar mechanisms might occur in insect vectors of plant viral pathogens but that they have been unnoticed because they cause no disease in their invertebrate hosts.

Wherever the above speculations may lead, it is useful to know that the previous work on LSNV can now be linked firmly to the sequence of WZSV 9 from China. This extends the known geographical distribution of LSNV to include China (Shi, et al., 2016) in addition to Thailand (Sritunyalucksana, et al., 2006), India (Kumar, et al., 2011; Prakasha, et al., 2007), Malaysia and Indonesia (Sittidilokratna, et al., 2009).

## Abbreviations lists

LSNV: Laem Singh virus
RT-PCR: Reverse-transcriptase polymerase chain reaction
RACE: Rapid amplification of cDNA ends

## Acknowledgements

We would like to thank the financial support from the Royal Society International Collaboration Awards 2019, the Global Challenges Research Fund (GCRF) to Prof. Grant D. Stentiford (Cefas/UK) and Dr. Kallaya Sritunyalucksana (BIOTEC, NSTDA/Thailand).

## References

Bell, T.A., Lightner, D.V., 1988. A handbook of normal shrimp histology. World Aquaculture Society, Baton Rouge, LA.

Kaur, N., Hasegawa, D.K., Ling, K.-S., Wintermantel, W.M., 2016. Application of genomics for understanding plant virus-insect vector interactions and insect vector control. Phytopathology. 106, 1213–1222.

Kumar, T.S., Krishnan, P., Makesh, M., Chaudhari, A., Purushothaman, C.S., Rajendran, K.V., 2011. Natural host-range and experimental transmission of Laem-Singh virus (LSNV). Dis. Aquat. Organ. 96, 21–27.

Lan, H., Wang, H., Chen, Q., Chen, H., Jia, D., Mao, Q., Wei, T., 2016. Small interfering RNA pathway modulates persistent infection of a plant virus in its insect vector. Sci. Rep. 6, 1–12.

Li, C.-X., Shi, M., Tian, J.-H., Lin, X.-D., Kang, Y.-J., Chen, L.-J., Qin, X.-C., Xu, J., Holmes, E.C., Zhang, Y.-Z., 2015. Unprecedented genomic diversity of RNA viruses in arthropods reveals the ancestry of negative-sense RNA viruses. eLife. 4, e05378.

Panphut, W., Senapin, S., Sriurairatana, S., Withyachumnarnkul, B., Flegel, T.W., 2011. A novel integrase-containing element may interact with Laem-Singh virus (LSNV) to cause slow growth in giant tiger shrimp. BMC Vet. Res. 7, 18.

Prakasha, B.K., Raju, R.P., Karunasagar, I., Karunasagar, I., 2007. Detection of Laem-Singh Virus (LSNV) in cultured *Penaeus monodon* from India. Dis. Aquat. Org. 77, 83–86.

Pratoomthai, B., Sakaew, W., Sriurairatana, S., Wongprasert, K., Withyachumnarnkul, B., 2008. Retinopathy in stunted black tiger shrimp *Penaeus monodon* and possible association with Laem-Singh virus (LSNV). Aquaculture. 284, 53–58.

Rubinstein, G., Czosnek, H., 1997. Long-term association of tomato yellow leaf curl virus with its whitefly vector Bemisia tabaci: effect on the insect transmission capacity, longevity and fecundity. J. Gen. Virol. 78, 2683–2689.

Shi, M., Lin, X.-D., Tian, J.-H., Chen, L.-J., Chen, X., Li, C.-X., Qin, X.-C., Li, J., Cao, J.-P., Eden, J.-S., 2016. Redefining the invertebrate RNA virosphere. Nature. 540, 539–543.

Sittidilokratna, N., Dangtip, S., Sritunyalucksana, K., Babu, R., Pradeep, B., C.V., M., Gudkovs, N., Walker, P.J., 2009 Detection of Laem-Singh virus in cultured *Penaeus monodon* shrimp from several sites in the Indo-Pacific region. Dis. Aquat. Org. 84, 195–200.

Sõmera, M., Sarmiento, C., Truve, E., 2015. Overview on Sobemoviruses and a proposal for the creation of the family *Sobemoviridae*. Viruses. 7, 3076–3115.

Sõmera, M., Truve, E., Eugenie, H., 2017. Create a new family of positive-sense RNA viruses, *Solemoviridae*. International Committee on Taxonomy of Viruses (ICTV).

Sritunyalucksana, K., Apisawetakan, S., Boon-nat, A., Withyachumnarnkul, B., Flegel, T.W., 2006. A new RNA virus found in black tiger shrimp *Penaeus monodon* from Thailand. Virus Res. 118, 31–38.

Tassetto, M., Kunitomi, M., Whitfield, Z.J., Dolan, P.T., Sánchez-Vargas, I., Garcia-Knight, M., Ribiero, I., Chen, T., Olson, K.E., Andino, R., 2019. Control of RNA viruses in mosquito cells through the acquisition of vDNA and endogenous viral elements. eLife. 8.

Whitfield, Z.J., Dolan, P.T., Kunitomi, M., Tassetto, M., Seetin, M.G., Oh, S., Heiner, C., Paxinos, E., Andino, R., 2017. The diversity, structure, and function of heritable adaptive immunity sequences in the *Aedes aegypti* genome. Curr. Biol. 27, 3511–3519. e3517.

