## Supplementary figures for "Full genome characterization of Laem Singh virus (LSNV) in shrimp *Penaeus monodon*"

**Figure S1.** Multiple sequence alignment of LSNV genome segment 1.

```
NC_033300_WSV9_S1  ATTGCGAAGAGCTGATTTAGGCGTTTGGTTCTCTGCAAGCTTCGGCCAATCACGTTCCCAATGGGTGTCTTTAAGAAGATCTTGTTGAAG 90
LSNV_RACE          ------------------------------------------------------------------------------------------ 1
LSNV-S1            ATTGCGAAGAGCTGATTTAGGCGTTCGGTTCTCTGCAAGCTTCGGCCAATCACGTTCCCAATGGGTGTCTTTAAGAAGATCTTGTTGAAG 90
Consensus          ATTGCGAAGAGCTGATTTAGGCGTTCGGTTCTCTGCAAGCTTCGGCCAATCACGTTCCCAATGGGTGTCTTTAAGAAGATCTTGTTGAAG 90

NC_033300_WSV9_S1  ACAATCAAGGAGGTGTTGGAAGAGGAGGAGTTGTCCACTCGTAAGAACCCCCTGACCATTGGGAACGTGGGTAAGTCTTGGTTGGCTGCT 180
LSNV_RACE          ------------------------------------------------------------------------------------------ 1
LSNV-S1            ACAATCAAGGAGGTGTTGGAAGAGGAGGAGTTGTCCACTCGTAAGAACCCCCTGACCATTGGGAACGTGGGTAAGTCTTGGTTGGCTGCT 180
Consensus          ACAATCAAGGAGGTGTTGGAAGAGGAGGAGTTGTCCACTCGTAAGAACCCCCTGACCATTGGGAACGTGGGTAAGTCTTGGTTGGCTGCT 180

NC_033300_WSV9_S1  GTTGGCACCTACAAGAAGCACCTTGTGGTTGGTGCTCTCGTGGGGGCAGGGGCCTTGGCTTACCGGTACTGGAAAAATAGAAAACGGCCC 270
LSNV_RACE          ------------------------------------------------------------------------------------------ 1
LSNV-S1            GTTGGCACCTACAAGAAGCACCTCGTGGTTGGTGCTCTCGTGGGGGCAGGGGCCTTGGCTTACCGGTACTGGAAAAATAGAAAACGGCCC 270
Consensus          GTTGGCACCTACAAGAAGCACCTCGTGGTTGGTGCTCTCGTGGGGGCAGGGGCCTTGGCTTACCGGTACTGGAAAAATAGAAAACGGCCC 270

NC_033300_WSV9_S1  TCGCGCGTAGTGAGGGACAGTTTCTTCAGGGCCGCTATAACAGGAGATGTGGTCCCGGAGAGTAAAGTGGCTGGTAGTCCAGAGGTGAAT 360
LSNV_RACE          ------------------------------------------------------------------------------------------ 1
LSNV-S1            TCGCGCGTAGTGAGGGACAGTTTCTTCAGGGCCGCTATAACAGGAGATGTGGTCCCGGAGAGTAAAGTGGCTGGTAGTCCAGAGGTGAAT 360
Consensus          TCGCGCGTAGTGAGGGACAGTTTCTTCAGGGCCGCTATAACAGGAGATGTGGTCCCGGAGAGTAAAGTGGCTGGTAGTCCAGAGGTGAAT 360

NC_033300_WSV9_S1  CTCTTGGCTCCCAATTCACAATGCCGAATTTTGTTGAAGAACTCGGATGGTGACTGGGTGTTGACTGGATGCGCTGTGCGCGTAGATCTG 450
LSNV_RACE          ------------------------------------------------------------------------------------------ 1
LSNV-S1            CTCTTGGCTCCCAATTCACAATGCCGAATTTTGTTGAAGAACTCGGATGGTGACTGGGTGTTGACTGGATGCGCTGTGCGCGTAGATCTG 450
Consensus          CTCTTGGCTCCCAATTCACAATGCCGAATTTTGTTGAAGAACTCGGATGGTGACTGGGTGTTGACTGGATGCGCTGTGCGCGTAGATCTG 450

NC_033300_WSV9_S1  CCCATTGGTCCCTTCCTGGTCTTTCCGCAACATTTGTGCGCTGGAGAGCTCG-CTGCTATGAGTTCTCGGGATGAGGTCATCCCCCTATC 539
LSNV_RACE          -----------------------------------------------GCTCGGCTGTAAGGAGTTCTCGGGATGAGGTCATCCCCTTATC 43
LSNV-S1            CCCATTGGTCCCTTCCTGGTCTTTCCGCAACATTTGTGCGCTGGAGAGCTCG-CTGCTATGAGTTCTCGGGATGAGGTCATCCCCCTATC 539
Consensus          CCCATTGGTCCCTTCCTGGTCTTTCCGCAACATTTGTGCGCTGGAGAGCTCGGCTGCTATGAGTTCTCGGGATGAGGTCATCCCCCTATC 540

NC_033300_WSV9_S1  GTTGGATGGCCTCGAGCCGATAGCGGCAGATTTGAGCATGCTCCCCTGTCCCTCTAAGCTGACCAGCACTGGACTCGCCTCGCCTACTAT 629
LSNV_RACE          GTTGGAATGCCTCGAGCCGATAGCGGGCAGATTGAGCATGCTCCCCTGTCCCTCTAAGCTGACCAGTACTGGACTCGTCTCGCCTACTAT 133
LSNV-S1            GTTGGATGGCCTCGAGCCGATAGCGGCAGATTTGAGCATGCTCCCCTGTCCCTCTAAGCTGACCAGCACTGGACTCGCCTCGCCTACTAT 629
Consensus          GTTGGATGGCCTCGAGCCGATAGCGGCAGATTTGAGCATGCTCCCCTGTCCCTCTAAGCTGACCAGCACTGGACTCGCCTCGCCTACTAT 630

NC_033300_WSV9_S1  AGGCGTTGTCCGCCCTGGGCAGCGTTGCGTCGTGAGTGTAGTCGGCGTTGATGGCTTGGGCACAACTGGACCTCTCCAGACTTCTCCCGT 719
LSNV_RACE          AGGCGTTGTCCGCCCTGGGCAGCGTTGCGTCGTGAGTGTAGTCGGCGTTGATGGCTTGGGCACAACTGGGCCTCTCCAGACTTCTCCCGT 223
LSNV-S1            AGGCGTTGTCCGCCCTGGGCAGCGTTGCGTTGTGAGTGTAGTCGGCGTTGATGGCTTGGGCACAACTGGGCCTCTCCAGACTTCTCCCGT 719
Consensus          AGGCGTTGTCCGCCCTGGGCAGCGTTGCGTCGTGAGTGTAGTCGGCGTTGATGGCTTGGGCACAACTGGGCCTCTCCAGACTTCTCCCGT 720

NC_033300_WSV9_S1  TTTCGGTAGGTTGATATATGGAGCAACCACTAATACCGGATATTCGGGGGCTGCCTATGCCTCCGCCAGGACTGTGTATGGAGTTCACAC 809
LSNV_RACE          TTTCGGTAGGTTGATATATGGAGCAACCACTAATACCGGATATTCGGGGGCTGCCTATGCCTCCGCCAGGACTGTGTATGGAGTTCACAC 313
LSNV-S1            TTTCGGTAGGTTGATATATGGAGCAACCACTAATACCGGATATTCGGGGGCTGCCTATGCCTCCGCCAGGACTGTGTATGGAGTTCACAC 809
Consensus          TTTCGGTAGGTTGATATATGGAGCAACCACTAATACCGGATATTCGGGGGCTGCCTATGCCTCCGCCAGGACTGTGTATGGAGTTCACAC 810

NC_033300_WSV9_S1  AAATGGCGGGAAACTCAACGGCGGTTTTGCTCTGTCGTATGTCTATGCCATGATGAAGGTGCTTAAAGGGGTCAGAGACGAGAGTTCAGA 899
LSNV_RACE          AAATGGCGGGAAACTCAACGGCGGTTTTGCTCTGTCGTATGTCTATGCCATGATGAAGGTGCTTAAAGGGGTCAGAGACGAGAGTTCAGA 403
LSNV-S1            AAATGGCGGGAAACTCAACGGCGGTTTTGCTCTGTCGTATGTCTATGCCATGATGAAGGTGCTTAAAGGGGTCAGAGACGAGAGTTCAGA 899
Consensus          AAATGGCGGGAAACTCAACGGCGGTTTTGCTCTGTCGTATGTCTATGCCATGATGAAGGTGCTTAAAGGGGTCAGAGACGAGAGTTCAGA 900

NC_033300_WSV9_S1  GAATTGGATTCATAGCGTCATGGCCGACAAGAGCACCCGGTTCTTGGAGATGGACTATGAGAGAGTAGCCGATAGCATAGTCATGCGTGG 989
LSNV_RACE          GAATTGGATTCATAGCGTCATGGCCGACAAGAGCACCCGGTTCTTGGAGATGGACTATGAGAGAGTAGCCGATAGCATAGTCATGCGTGG 493
LSNV-S1            GAACTGGATTCATAGCGTCATGGCTGACAAGAGCACCCGGTTCTTGGAGATGGACTATGAGAGAGTAGCCGATAGCATAGTCATGCGTGG 989
Consensus          GAATTGGATTCATAGCGTCATGGCCGACAAGAGCACCCGGTTCTTGGAGATGGACTATGAGAGAGTAGCCGATAGCATAGTCATGCGTGG 990

NC_033300_WSV9_S1  TGATGATGGCTTCTATCACGCTATCTCAGGCCCTGCAGCCGAGGGCTTGGTGAAGAAGCTTGATAAGTACAAGTATGATGCTGGCATGGT 1079
LSNV_RACE          TGATGATGGCTTCTATCACGCTATCTCAGGCCCTGCAGCTGAGGGCTTGGTGAAGAAGCTTGATAAGTACAAGTATGATGCTGGCATGGT 583
LSNV-S1            TGATGATGGCTTCTATCACGCTATCTCAGGCCCTGCAGCTGAGGGCTTGGTGAAGAAGCTTGATAAGTACAAGTATGATGCTGGCATGGT 1079
Consensus          TGATGATGGCTTCTATCACGCTATCTCAGGCCCTGCAGCTGAGGGCTTGGTGAAGAAGCTTGATAAGTACAAGTATGATGCTGGCATGGT 1080

NC_033300_WSV9_S1  AGATTTCCGGTCTAAGACTGGGCATACTTGGTCCGATATTGATCCCAATCATTGGGAGTCAGCAAACCCTGAGACTGCACAAGTACTCGC 1169
LSNV_RACE          AGATTTCCGGTCTAAGACTGGGCATACTTGGTCCGATATTGATCCCAATCATTGGGAGTCAGCAAACCCTGAGACTGCGCAAGTACTCGC 673
LSNV-S1            AGATTTCCGGTCTAAGACTGGGCATACTTGGTCCGATATTGATCCCAATCATTGGGAGTCAGCAAACCCTGAGACAGCACAAGTACTCGC 1169
Consensus          AGATTTCCGGTCTAAGACTGGGCATACTTGGTCCGATATTGATCCCAATCATTGGGAGTCAGCAAACCCTGAGACTGCACAAGTACTCGC 1170

LSNV-S1            AGATTTCCGGTCTAAGACTGGGCATACTTGGTCCGATATTGATCCCAATCATTGGGAGTCAGCAAACCCTGAGACAGCACAAGTACTCGC 1169
Consensus          AGATTTCCGGTCTAAGACTGGGCATACTTGGTCCGATATTGATCCCAATCATTGGGAGTCAGCAAACCCTGAGACTGCACAAGTACTCGC 1170

NC_033300_WSV9_S1  TTCGGCAGATACAGTGCCCCAGGGTTTTCACTTGGACCCAAGCTCCCGGGAGTCCCCGTACATAGAGGAATTAGCAAGCCAAATCCGAAT 1259
LSNV_RACE          TTCGGCAGATACAGTGCCCCAGGGTTTTCACTTGGACCCAAGCTCCCGGGAGTCCCCGTACATAGAGGAATTAGCAAGCCAAATCCGAAT 763
LSNV-S1            TTCGGCAGATACAGTGCCCCAGGGTTTTCACTTGGACCCAAGCTCCCGGGAATCCCCGTACATAGAGGAATTAGCAAGCCAAATCCGAAT 1259
Consensus          TTCGGCAGATACAGTGCCCCAGGGTTTTCACTTGGACCCAAGCTCCCGGGAGTCCCCGTACATAGAGGAATTAGCAAGCCAAATCCGAAT 1260

NC_033300_WSV9_S1  GTTCGAGGACAGATTGAAAAGTTCTTTGGTGGACACTGCCTTAAAGAAGCAGATAATGGATACGTTCCAGCAGACACTGGAACCCTCGCG 1349
LSNV_RACE          GTTCGAGGACAGATTGAAAAGTTCTTTGGTGGACACTGCCTTAAAGAAGCAGATAATGGATACGTTCCAGCAGACACTGGAACCCTCGCG 853
LSNV-S1            GTTCGAGGACAGATTGAAAAGTTCTTTGGTGGACACTGCCTTAAAGAAGCAGATAATGGATACGTTCCAGCAGACACTGGAACCTTCGCG 1349
Consensus          GTTCGAGGACAGATTGAAAAGTTCTTTGGTGGACACTGCCTTAAAGAAGCAGATAATGGATACGTTCCAGCAGACACTGGAACCCTCGCG 1350

NC_033300_WSV9_S1  GTCTCTCACTCTTTCGCTCTGCAGGCCAGAACTGCAGCACAGGCTCGCGAAAGTTATGTTCGGCCTTCATTAGAGGAAGTGGCCTGGGTT 1439
LSNV_RACE          GTCTCTCACTCTTTCGCTCTGCAGGCCAGAACTGCAGCACAGGCTCGCGAAAGTTATGTTCGGCCTTCATTAGAGGAAGTGGCCTGGGTT 943
LSNV-S1            GTCTCTCACTCTTTCGCTCTGCAGGCCAGAACTGCAGCACAGGCTCGCGAAAGTTATGTTCGGCCTTCATTAGAGGAAGTGGCCTGGGTT 1439
Consensus          GTCTCTCACTCTTTCGCTCTGCAGGCCAGAACTGCAGCACAGGCTCGCGAAAGTTATGTTCGGCCTTCATTAGAGGAAGTGGCCTGGGTT 1440

NC_033300_WSV9_S1  TTGCATCGGATGGAGGAAGCTTATGCTGCCGTCCGGTGGGATATACCCAGTGACTTCCTCTCTCGCGAACATTATGAGAGAACCCTGCTG 1529
LSNV_RACE          TTGCATCGGATGGAGGAAGCTTATGCTGCCGTCCGGTGGGATATACCCAGTGACTTCCTCTCTCGCGAACATTATGAGAGAACCCTGCTG 1033
LSNV-S1            TTGCATCGGATGGAGGAAGCTTATGCTGCCGTCCGGTGGGATATACCCAGTGACTTCCTCTCTCGCGAACATTATGAGAGAACCCTGCTG 1529
Consensus          TTGCATCGGATGGAGGAAGCTTATGCTGCCGTCCGGTGGGATATACCCAGTGACTTCCTCTCTCGCGAACATTATGAGAGAACCCTGCTG 1530
```

```
NC_033300_WSV9_S1  CGTCTTGATTTGCAAGCTTCCCCAGGTTATCCGTATTTGCGTGAGGCTGCCACCATTGGTCTGTGGCTTGGTGTGACCCTTGAGGGTCAA 1619
LSNV_RACE          CGTCTTGATTTGCAAGCTTCCCCAGGTTATCCGTATTTGCGTGAGGCTGCCACCATTGGTCTGTGGCTTGGTGTGACCCTTGAGGGTCAA 1123
LSNV-S1            CGTCTTGATTTGCAAGCTTCCCCAGGTTATCCGTATTTGCGTGAGGCTGCCACCATTGGTCTGTGGCTTGGTGTGACCCTTGAGGGTCAA 1619
Consensus          CGTCTTGATTTGCAAGCTTCCCCAGGTTATCCGTATTTGCGTGAGGCTGCCACCATTGGTCTGTGGCTTGGTGTGACCCTTGAGGGTCAA 1620

NC_033300_WSV9_S1  TTTGCCCCGGACCAAGTTGAGCGGCTTTGGCAACATATCCTTCTTATTTTGTCCGGTGAGTTTGATCATTACTACAGAGTGTTTGTTAAG 1709
LSNV_RACE          TTTGCCCCGGACCAAGTTGAGCGGCTTTGGCAACATATCCTTCTTATTTTGTCCGGTGAGTTTGATCATTACTACAGAGTGTTTGTTAAG 1213
LSNV-S1            TTTGCCCCGGACCAAGTTGAGCGGCTTTGGCAACATATCCTTCTCATTTTGTCCGGTGAGTTTGATCATTACTATAGAGTGTTTGTTAAG 1709
Consensus          TTTGCCCCGGACCAAGTTGAGCGGCTTTGGCAACATATCCTTCTTATTTTGTCCGGTGAGTTTGATCATTACTACAGAGTGTTTGTTAAG 1710

NC_033300_WSV9_S1  GATGAAGTTCACAGGAAGAAGAAGGCGGATGAGGGGAGGTGGCGACTGATATTAGCATCTGCATTGCCCATGCAAGTTCTGTGGCACATG 1799
LSNV_RACE          GATGAAGTTCACAGGAAGAAGAAGGCGGATGAGGGGAGGTGGCGACTGATATTAGCATCTGCATTGCCCATGCAAGTTTTGTGGCATATG 1303
LSNV-S1            GATGAAGTTCACAGGAAGAAGAAGGCGGATGAGGGGAGGTGGCGACTGATATTAGCATCTGCATTGCCCATGCAAGTTCTGTGGCATATG 1799
Consensus          GATGAAGTTCACAGGAAGAAGAAGGCGGATGAGGGGAGGTGGCGACTGATATTAGCATCTGCATTGCCCATGCAAGTTCTGTGGCATATG 1800

NC_033300_WSV9_S1  CTCTTTGCGCCTATGAATGACTTGGAAGCAGAGAAGATCTTCCACACACCTTCAGCATTTGGAGTCTCATTTGTTTATGGTGAGTGGAAG 1889
LSNV_RACE          CTCTTTGCGCCTATGAATGACTTGGAAGCAGAGAAGATCTTCCACACACCTTCAGCATTTGGAGTCTCATTTGTTTATGGTGAGTGGAAG 1393
LSNV-S1            CTCTTTGCGCCTATGAATGACTTGGAAGCAGAGAAGATCTTCCACACACCTTCAGCATTTGGAGTCTCATTTGTTTATGGTGAGTGGAAG 1889
Consensus          CTCTTTGCGCCTATGAATGACTTGGAAGCAGAGAAGATCTTCCACACACCTTCAGCATTTGGAGTCTCATTTGTTTATGGTGAGTGGAAG 1890

NC_033300_WSV9_S1  ATGTTCAAGAACTACTGCAGAGTCAAACGCTTAGAGGTTGCCATAGACAAGTCTGGTTGGGATTGGAACGCCCCTGGGTGGGTGTTTGAT 1979
LSNV_RACE          ATGTTCAAGAATTACTGCAAAGTCAAACGCTTAGAGGTTGCCATAGACAAGTCTGGTTGGGATTGGAACGCCCCTGGGTGGGTGTTTGAT 1483
LSNV-S1            ATGTTCAAGAATTACTGCAAAGTCAAACGCTTAGAGGTTGCCATAGACAAGTCTGGTTGGGATTGGAACGCCCCTGGGTGGGTGTTTGAT 1979
Consensus          ATGTTCAAGAATTACTGCAAAGTCAAACGCTTAGAGGTTGCCATAGACAAGTCTGGTTGGGATTGGAACGCCCCTGGGTGGGTGTTTGAT 1980

NC_033300_WSV9_S1  GCTGATCTGCAGTTGCGCTACCGCCTCTGTAACCAGGCTGAGCGGCCTGCAGGACATTTGTGGTTTGCTCTAGCCAGGAAGCTGTATGAT 2069
LSNV_RACE          GCTGATTTGCAGTTGCGCTACCGCCTCTGTAACCAGGCTGAGCGGCCTGCAGGACATTTGTGGTTTGCTCTAGCCAGGAAGCTGTATGAT 1573
LSNV-S1            GCTGATTTGCAGTTGCGCTACCGCCTCTGTAACCAGGCTGAGCGGCCTGCAGGACATTTGTGGTTTGCTCTAGCCAGGAAGCTGTATGAT 2069
Consensus          GCTGATTTGCAGTTGCGCTACCGCCTCTGTAACCAGGCTGAGCGGCCTGCAGGACATTTGTGGTTTGCTCTAGCCAGGAAGCTGTATGAT 2070

NC_033300_WSV9_S1  GATGCCTTTGAACACTCCCGTTGCCTTCTCCCGAGTGGTCAGGTTTACGTGCAAGAGTTCTCAGGCTTCATGAAGTCAGGCATTGTGAAT 2159
LSNV_RACE          GATGCCTTTGAACACTCCCGTTGCCTTCTCCCGAGTGGTCAGGTTTACGCGCAAGAGTTCTCAGGCTTCATGAAGTCAGGCATTGTGAAT 1663
LSNV-S1            GATGCCTTTGAACACTCCCGTTGCCTTCTCCCGAGTGGTCAGGTTTACGTGCAAGAGTTCTCAGGCTTCATGAAGTCAGGCATTGTGAAT 2159
Consensus          GATGCCTTTGAACACTCCCGTTGCCTTCTCCCGAGTGGTCAGGTTTACGTGCAAGAGTTCTCAGGCTTCATGAAGTCAGGCATTGTGAAT 2160

NC_033300_WSV9_S1  ACTATCTCCACCAATTCTCATGCCCAGATCATGCTGCATATGCTTGCTTGCAAGCGCTCGGGTGAGCCCGTGACTCCTATATTAGCCTGC 2249
LSNV_RACE          ACTATCTCCACCAATTCTCATGCCCAGATCATGCTGCATATGCTTGCTTGCAAGCGCTCGGGTGAGCCCGTGACTCCTATATTAGCCTGC 1753
LSNV-S1            ACTATCTCCACCAATTCTCATGCTCAGATCATGCTGCACATGCTTGCTTGCAAGCGCTCGGGTGAGCCCGTGACTCCTATATTAGCCTGC 2249
Consensus          ACTATCTCCACCAATTCTCATGCCCAGATCATGCTGCATATGCTTGCTTGCAAGCGCTCGGGTGAGCCCGTGACTCCTATATTAGCCTGC 2250

NC_033300_WSV9_S1  GGTGATGACACTATTCAAGCAGCTACCTCAGCCGGTTATATTGAAGAGCTGGCGAAGGCAGGGTGCATTGTTAAGAGTGTCGATCGCAAG 2339
LSNV_RACE          GGTGATGACACTATTCAAGCAGCTACCTCAGCCGGTTATATTGAAGAGCTGGCGAAGGCAGGGTGCATTGTTAAGAGTGTCGATCGCAAG 1843
LSNV-S1            GGTGATGACACTATTCAAGCAGCTACCTCAGCCGGTTATATTGAAGAGCTGGCGAGGGCAGGGTGCATTGTTAAGAGTGTCGATCGCAAG 2339
Consensus          GGTGATGACACTATTCAAGCAGCTACCTCAGCCGGTTATATTGAAGAGCTGGCGAAGGCAGGGTGCATTGTTAAGAGTGTCGATCGCAAG 2340

NC_033300_WSV9_S1  CTAGAGTTTATGGGCTTTAATTTTGAGGGCGCCATGCAACCAATCTACACCGTGAAACACATCGCATCTTTTACTTATAAGAGCGAGGAG 2429
LSNV_RACE          CTAGAGTTTATGGGTTTTAACTTTGAGGGCGCCATGCAACCAATCTACACCATGAAACACATCGCATCTTTTACTTATAAGAGTGAGGAG 1933
LSNV-S1            CTAGAGTTTATGGGTTTTAACTTTGAGGGCGCCATGCAACCAATCTACACCATGAAACACATCGCATCTTTCACTTATAAGAGTGAGGAG 2429
Consensus          CTAGAGTTTATGGGTTTTAACTTTGAGGGCGCCATGCAACCAATCTACACCATGAAACACATCGCATCTTTTACTTATAAGAGTGAGGAG 2430

NC_033300_WSV9_S1  CTTCATGGTGAGATTTTAGATAGCATGTGCAGAATGTACGCTCACCACCCCTGGTTTGACTGTTGGAAGATGTTAGCAGGCTCTTTGGGT 2519
LSNV_RACE          CTTCATGGTGAGATTTTAGATAGCATGTGCAGAATGTACGCTCACCACCCCTGGTTTGACTGTTGGAAGATGTTAGCAGGCTCTTTGGGC 2023
LSNV-S1            CTTCATGGTGAGATTTTAGATAGCATGTGCAGAATGTACGCTCACCACCCCTGGTTTGACTGTTGGAAGATGTTAGCAGGCTCTTTGGGC 2519
Consensus          CTTCATGGTGAGATTTTAGATAGCATGTGCAGAATGTACGCTCACCACCCCTGGTTTGACTGTTGGAAGATGTTAGCAGGCTCTTTGGGC 2520

NC_033300_WSV9_S1  CATAATATGAAGTCTAGAGCCTGGTATCAGTTCTTCCTTGATAGTACAGCCAATATCAGGATCACGAAATCCATTTAGATAGAGGATGAC 2609
LSNV_RACE          CATAATATGAAGTCTAGAGCCTGGTATCAGTTCTTCCTTGATAGTACAGCCAATATCAGGATCACGAAGTCCATTTAGATAGAGGATGAC 2113
LSNV-S1            CATAATATGAAGTCTAGAGCCTGGTATCAGTTCTTCCTTGATAGTACAGCCAATATCAGGATCACGAAGTCCATTTAGATAGAGGATGAC 2609
Consensus          CATAATATGAAGTCTAGAGCCTGGTATCAGTTCTTCCTTGATAGTACAGCCAATATCAGGATCACGAAGTCCATTTAGATAGAGGATGAC 2610

NC_033300_WSV9_S1  GTGAGCTGGCTATGTCTTCTACCTTTCCTGTAACGTGGCGGTTCTGAGGGGGCGTGTGCCAACACGTTTCTGGGGC-------------- 2685
LSNV_RACE          GTGAGCTGGCTATGTTTTCTACCTTTCCTGTAACGTGGCGGTTCTGAGGGGGCGTGTGCCAATACGTTTCTGGGGCAAAAAAAAAAAAAA 2203
LSNV-S1            GTGAGCTGGCTATGTTTTCTACCTTTCCTGTAACGTGGCGGTTCTGAGGGGGCGTGTGCCAACACGTTTCTGGGG--------------- 2684
Consensus          GTGAGCTGGCTATGTTTTCTACCTTTCCTGTAACGTGGCGGTTCTGAGGGGGCGTGTGCCAACACGTTTCTGGGGCAAAAAAAAAAAAAA 2700

NC_033300_WSV9_S1  --- 2685
LSNV_RACE          AAA 2206
LSNV-S1            --- 2684
Consensus          AAA 2703
```

**Figure S2.** Multiple sequence alignment of LSNV genome segment 2.

```
NC_033295_WSV9_S2  CGGGAATATTAGAAAAGGTTTGGTCTAACATTTCAAGCAGACCCAGAACAACATGACTAACCAGCATGTTTGTCCAATCTGCAAGCGTTC 90
LSNV-S2            CGGGAATATTAGAAAAGGTTTGGTCTAACATTTCAAGCAGACCCAGAACAACATGACTAACCAGCATGTTTGTCCAATCTGCAAGCGTTC 90
Consensus          CGGGAATATTAGAAAAGGTTTGGTCTAACATTTCAAGCAGACCCAGAACAACATGACTAACCAGCATGTTTGTCCAATCTGCAAGCGTTC 90

NC_033295_WSV9_S2  TTTCAGAACAGCCCAAGGGTTGGCGGACCACAAACGTGACCGGCATGAAGCTTGTTCCACCGGACCTGCCCCTAAGCCTCAAAGGGTCAG 180
LSNV-S2            TTTCAGAACAGCCCAAGGGTTGGCGGACCACAAACGTGACCGGCATGAAGCTTGTTCCACCGGACCTGCCCCTCAACCTAAAAGGGTCAG 180
Consensus          TTTCAGAACAGCCCAAGGGTTGGCGGACCACAAACGTGACCGGCATGAAGCTTGTTCCACCGGACCTGCCCCTAAACCTAAAAGGGTCAG 180

NC_033295_WSV9_S2  GCCTAAGAGGGAACAGCGTCAGGCCGCTCCCGTTAGTGTGCCCACAAGTTCTGGGCCCGCTCGTTACTCCAATCGAGTGACTTTGCAGGG 270
LSNV-S2            GCCTAGGAGGGAACAGCGTCAGGCCGCTCCCGTTAGTGTGCCCACAAGTTCTGGGCCCGCTCGTTACTCCAATCGAGTGACTTTGCAGGG 270
Consensus          GCCTAAGAGGGAACAGCGTCAGGCCGCTCCCGTTAGTGTGCCCACAAGTTCTGGGCCCGCTCGTTACTCCAATCGAGTGACTTTGCAGGG 270

NC_033295_WSV9_S2  GAGCAACGAGCGGTTCCGGACCGACGATTATCCTGCGTCCAGACTGAAGACGGGGACTTTGCTCACAACGATTCCTGTGAGCCCAGACCT 360
LSNV-S2            GAGCAACGAGCGGTTCCGGACCGACGATTATCCTGCGTCCAGACTGAAGACGGGGACTTTGCTCACAACAATTCCTGTGAGCCCAGACCT 360
Consensus          GAGCAACGAGCGGTTCCGGACCGACGATTATCCTGCGTCCAGACTGAAGACGGGGACTTTGCTCACAACAATTCCTGTGAGCCCAGACCT 360

NC_033295_WSV9_S2  GGTCCCCAGGCTTTCCACGCAGGCTAAGGCGTATCAGCGCATCAAGTATCATTCTGTTGCGCTGCATATCGACGCCAAGGGTTCGACCGC 450
LSNV-S2            GGTCCCCAGGCTTTCCACGCAGGCTAAGGCGTATCAGCGCATTAAGTATCATTCTGTTGCGCTGCATATCGACGCCAAGGGTTCGACCGC 450
Consensus          GGTCCCCAGGCTTTCCACGCAGGCTAAGGCGTATCAGCGCATCAAGTATCATTCTGTTGCGCTGCATATCGACGCCAAGGGTTCGACCGC 450

NC_033295_WSV9_S2  CCAGTCGGGCGGCTACTTGGCTGTGTTCATCAGCGATATCACTGACGAAGAGATTTCGCTGGAGCGAGCCGGTTCCTTTGGCGGTAGTGT 540
LSNV-S2            CCAGTCGGGCGGCTACTTGGCTGTGTTCATCAGCGATATCACTGACGAAGAGATTTCGCTGGAGCGAGCCGGTTCCTTTGGCGGTAGTGT 540
Consensus          CCAGTCGGGCGGCTACTTGGCTGTGTTCATCAGCGATATCACTGACGAAGAGATTTCGCTGGAGCGAGCCGGTTCCTTTGGCGGTAGTGT 540

NC_033295_WSV9_S2  CGGCAAGAAGTATTGGGAGAATGCCACGGTGACCGCGACCAATTTGACGCCTCTCTTCTACACTAGTAGGGGAGAGGATGAGCGGTTGTG 630
LSNV-S2            CGGCAAGAAGTATTGGGAGAATGCCACGGTGACCGCGACCAATCTGACGCCTCTCTTCTACACCAGTAGGGGAGAGGATGAGCGGTTGTG 630
Consensus          CGGCAAGAAGTATTGGGAGAATGCCACGGTGACCGCGACCAATCTGACGCCTCTCTTCTACACCAGTAGGGGAGAGGATGAGCGGTTGTG 630

NC_033295_WSV9_S2  GTCTCCTGGCTATTTCGCAATTTATTGTGATGGTTCTTCCAATACTGCCATCTCATTGACGTATCGGTTGACTTGGAGTGTGACCTTGTC 720
LSNV-S2            GTCTCCTGGCTATTTCGCAATTTATTGTGATGGTTCTTCCAATACTGCTATCTCATTGACGTATCGGTTGACTTGGAGTGTGACCTTGTC 720
Consensus          GTCTCCTGGCTATTTCGCAATTTATTGTGATGGTTCTTCCAATACTGCCATCTCATTGACGTATCGGTTGACTTGGAGTGTGACCTTGTC 720

NC_033295_WSV9_S2  GGTGCCTTCTTCCGAGTTGAAGCAGGCTGATGTCATCACGACCATGGTGGATCTGTGGCCGACTGCTGGCAAGGCTAGCTTTTCGGACAA 810
LSNV-S2            GGTGCCTTCTTCCGAGTTGAAGCAGGCTGATGTCATCACGACCATGGTGGATCTGTGGCCGACTGCTGGCAAGGCTAGCTTTTCGGACAA 810
Consensus          GGTGCCTTCTTCCGAGTTGAAGCAGGCTGATGTCATCACGACCATGGTGGATCTGTGGCCGACTGCTGGCAAGGCTAGCTTTTCGGACAA 810

NC_033295_WSV9_S2  GGATGGCTCTGAAGAGAATCTCTTTGAGCCCTCCTTGCCAGTTGGCACTGTGGTGAGGTTTCCTTATGCAGTTGGCATAGAGTACAAGGA 900
LSNV-S2            GGATGGCTCTGAAGAGAATCTCTTTGAGCCCTCCTTGCCAGTTGGCACTGTGGTGAGGTTTCCTTATGCAGTTGGCATAGAGTACAAGGA 900
Consensus          GGATGGCTCTGAAGAGAATCTCTTTGAGCCCTCCTTGCCAGTTGGCACTGTGGTGAGGTTTCCTTATGCAGTTGGCATAGAGTACAAGGA 900

NC_033295_WSV9_S2  GGGGGCTGGTGACACCGGAACTGTCCCTATTATGTTCGCCGTGGTGACAGCCTCGTCAGGGCTTAAGGCCAGTAATGACAGGGTTGACAC 990
LSNV-S2            GGGGGCTGGTGACACCGGAACTGTCCCTATTATGTTCGCCGTGGTGACAGCCTCGTCAGGGCTTAAGGCCAGTAATGACAGGGTTGACAC 990
Consensus          GGGGGCTGGTGACACCGGAACTGTCCCTATTATGTTCGCCGTGGTGACAGCCTCGTCAGGGCTTAAGGCCAGTAATGACAGGGTTGACAC 990

NC_033295_WSV9_S2  CTCGGTGGTTTGGCAGTCCGATGTTTCCCGTCGACTATTGTACCCCAAGTCCATCAGGTTGATTGTCGAGAAGCGTCCGGGGGAAGTGAG 1080
LSNV-S2            CTCGGTGGTTTGGCAGTCCGATGTTTCCCGTCGACTATTGTACCCCAAGTCCATCAGGTTGATTGTCGAGAAGCGTCCGGGGGAAGTGAG 1080
Consensus          CTCGGTGGTTTGGCAGTCCGATGTTTCCCGTCGACTATTGTACCCCAAGTCCATCAGGTTGATTGTCGAGAAGCGTCCGGGGGAAGTGAG 1080

NC_033295_WSV9_S2  CCGGGCGGGCCAGGTGAGCCAGTTTTCGCCTTGTCCGCCCACCAGTTTAGCTTCATCCGAGACGCCTTCCACCGAGACCGTAGGGAACAA 1170
LSNV-S2            CCGGGCGGGCCAGGTGAGCCAGTTTTCGCCTTGTCCGCCCACCAGTTTAGCTTCATCCGAGACGCCTTTCACCGAGACCGTAGGGAACAA 1170
Consensus          CCGGGCGGGCCAGGTGAGCCAGTTTTCGCCTTGTCCGCCCACCAGTTTAGCTTCATCCGAGACGCCTTCCACCGAGACCGTAGGGAACAA 1170

NC_033295_WSV9_S2  CGAAAACAGACCCACGTGCTCGGTGAGATTTACCACGCCCTCGTCAAGCGACTTGATGAAATCGATGCACGGTTGCAAGACCTCGATGAG 1260
LSNV-S2            CGAAAACAGACCCACGTGCTCGGTGAGATTTACCACGCCCTCGTCAAGCGACTTGATGAAATCGACGCACGGTTGCAAGACCTCGATGAG 1260
Consensus          CGAAAACAGACCCACGTGCTCGGTGAGATTTACCACGCCCTCGTCAAGCGACTTGATGAAATCGACGCACGGTTGCAAGACCTCGATGAG 1260

NC_033295_WSV9_S2  CGACTTGAGGGCAAGGCCGGTTTGCACGGTTTCATCGACATTGCCTCCCTCCCCAAACCCGTCGCAGGAGTCTATTCTATTGACTTTTCT 1350
LSNV-S2            CGACTTGAGGGCAAGGCCGGTTTGCACGGTTTCATCGACATTGCCTCCCTCCCCAAACCCGTCGCAGGAGTCTACTCTATTGACTTTTCT 1350
Consensus          CGACTTGAGGGCAAGGCCGGTTTGCACGGTTTCATCGACATTGCCTCCCTCCCCAAACCCGTCGCAGGAGTCTACTCTATTGACTTTTCT 1350

NC_033295_WSV9_S2  GGAGAGCCAGGTCCCTCGCTATCAGGCTGCACGGATTGTGAGTCAAGAGGTGAAGACTCATGAGTTTCCCCTCTTGCATCATCGCAATCA 1440
LSNV-S2            GGAGAGCCAGGTCCCTCGCTATCAGGCTGCACGGATTGTGAGTCAAGAGGTGAAGACTCATGAGTTTCCCCTCTTGCATCATCGCAATCA 1440
Consensus          GGAGAGCCAGGTCCCTCGCTATCAGGCTGCACGGATTGTGAGTCAAGAGGTGAAGACTCATGAGTTTCCCCTCTTGCATCATCGCAATCA 1440

NC_033295_WSV9_S2  TTTGCGGCTTGAGATGCATTGGGATCTGGTTCCTTTGATGAGGGGGAGGCGAATTCCGGTGTACGAGCTGAGTAGGCATTTTGATGATGA 1530
LSNV-S2            TTTGCGGCTTGAGATGCATTGGGATCTGGTTCCTTTGATGAGGGGGAGGCGAATTCCGGTGTACGAGCTGAGTAGGCATTTTGATGATGA 1530
Consensus          TTTGCGGCTTGAGATGCATTGGGATCTGGTTCCTTTGATGAGGGGGAGGCGAATTCCGGTGTACGAGCTGAGTAGGCATTTTGATGATGA 1530

NC_033295_WSV9_S2  CTTCTCCTTGAAGTTCACTATCAACAATACCGTCAGCTTCCGAGAAGTTGTCGAGGCAGTGGAGGTTTTGAACAGAGGTAGGTACTTCTG 1620
LSNV-S2            CTTCTCTTTGAAGTTCACTATCAATTATACCGTCAGCTTCCGAGAAGTTGTCGAGGCAGTGGAGGTTTTGAACAGAGGTAGGTACTTCTG 1620
Consensus          CTTCTCCTTGAAGTTCACTATCAACAATACCGTCAGCTTCCGAGAAGTTGTCGAGGCAGTGGAGGTTTTGAACAGAGGTAGGTACTTCTG 1620

NC_033295_WSV9_S2  TTTGGGCTTTATAGCACTGTGCCTGCAACATCATTTTGGTGCAGACTCTAGTGAAGTAGTGCCTGAGGAGTTCCCGTCGGTGTGGTACGA 1710
LSNV-S2            TTTGGGCTTTATAGCACTGTGCCTGCAACATCATTTTGGTGCAGACTCTAGTGAAGTAGTGCCTGAGGAGTTCCCGTCGGTGTGGTACGA 1710
Consensus          TTTGGGCTTTATAGCACTGTGCCTGCAACATCATTTTGGTGCAGACTCTAGTGAAGTAGTGCCTGAGGAGTTCCCGTCGGTGTGGTACGA 1710

NC_033295_WSV9_S2  TGAGAGCTTCCATTACCGCCTGCCTTGCCGATCCTGCCATGATAACATGGAATCTGACTACTCTTTTGAGGAGTTAGACCCGGAGGTGGA 1800
LSNV-S2            TGAGAGCTTCCATTACCGCCTGCCTTGCCGATCCTGCCATGATAACATGGAATCTGACTACTCTTTTGAGGAGTTAGACCCGGAGGTGGA 1800
Consensus          TGAGAGCTTCCATTACCGCCTGCCTTGCCGATCCTGCCATGATAACATGGAATCTGACTACTCTTTTGAGGAGTTAGACCCGGAGGTGGA 1800

NC_033295_WSV9_S2  GTATGATTAGTTCAGCTCCAAATGACTCGGCGTGAGGTCATGTGGCACGCCTCCCCAGAACCGCCACATGCGTCAGAAGGAAAACTGACG 1890
LSNV-S2            GTGTGATTAGTTCAGCTCCAAATGACTCGGCGTGAGGTCATGTGGCA------------------------------------------- 1847
Consensus          GTATGATTAGTTCAGCTCCAAATGACTCGGCGTGAGGTCATGTGGCACGCCTCCCCAGAACCGCCACATGCGTCAGAAGGAAAACTGACG 1890

NC_033295_WSV9_S2  TAAAACAAACCCTTGAGGAGCTGTCGCCAGCCAG 1924
LSNV-S2            ---------------------------------- 1847
Consensus          TAAAACAAACCCTTGAGGAGCTGTCGCCAGCCAG 1924
```

**Figure S3.** Multiple sequence alignment of deduced amino acids corresponding to predicted ORF1 of LSNV genome segment 1. Red-labelled letters indicate more than 60% amino acids identity.

```
YP_009337868      ------------------------------------------------------------MGVFKKILLKTIKEVLEEEELSTRKNPLTI 30
LSNVS1_PCR_ORF1   ------------------------------------------------------------MGVFKKILLKTIKEVLEEEELSTRKNPLTI 30
ASM94003          -----------------------------------------ERLFESMSVTTFLNNLWESASEPAPLTLEQVREQFPHLFEQRPSSAFLR 49
YP_009330079      --------------------------------------------------------------------------------MSWKDYLLVS 10
YP_009329959      --------------------------------------------------------MTTASFMFALARKQLPTAPPNPGWGIGLARPTLR 34
AMO03214          ------------------------------------------------------------------------------------------ 1
QED21501          ---------------------------------------------------MDTLIDTINVHRDLMIPIPGWKDLVLSTYREMPRWGVLV 39
AWA82253          MAAEAFLKIIPALSTGLVAGVGALGYHFSNEMAALRADAAAMAASLAAAKASAAAKTSIVEGVWKVVRGAGEITCTAIKESDLRTKILVG 90
AMO03212          -------------------------------------------------------------------------------------YALIA 5
QED21515          ---------------------------------------------------MENVNCAVSACVARTPSRT--KERLFKVLEVATSRPVLL 37
YP_009337259      ---------------------------------------------------MLSMMRNSVQVVPMAASALAGAAAYQVGKIASQAYDSLK 39

YP_009337868      GNVGKSWLAAVGTYKKHL------VVGALVGAGALAYRYWKNRKRPSRVVRDSFFRAA-------------ITGDVVPESKVAGSPEVNL 101
LSNVS1_PCR_ORF1   GNVGKSWLAAVGTYKKHL------VVGALVGAGALAYRYWKNRKRPSRVVRDSFFRAA-------------ITGDVVPESKVAGSPEVNL 101
ASM94003          AAAGRAVETTSGFFKGTVPETVPLKWVLGAGALAVAAYYVKDPLCKGVVSTSSALSAVCRSLFGKVKIRSVPLNDYLAESVVAGSQEMSM 139
YP_009330079      GGVYGLWMGHKIGVGS--------KISAVAGMVIPGFRKLKWAVGRAEVVSQPAAS-----------------NHLRLESRKDGSEEVSL 75
YP_009329959      EGLVLVGGVAATAASG-------YAVYSVGCWTMSACRRLRSWWAKPLKLRPAPSGPVVG--------------ECVPESAVPGSEERPM 103
AMO03214          ----------------------------QAGKLVPGLRTVKAWFGRIEVVLDKTVHS---------------KSSCILESRRAGSDEVDL 47
QED21501          G--GIVGGVVACKGVR--------PTGRLIGRMFGWTPVVARSVLTACGVPPPAVPSPK------------TDLQNTFESIREGSVEKPM 107
AWA82253          TAAYAAWLTYEYVPRP--------KVSKVAGAVLPGARRVRNWFGKAEIVADPRILRG-------------NQNSMLAESRRDGSAEQDL 159
AMO03212          GASYVTWHLTKEGCVG--------WIAKQAGKLVPGFRTIRNIVGKIEVVPGDGLTR----------------NRYVMESRKAGSDEIGM 71
QED21515          TAAGVAGLCMIGRHTG--------IHSKIIGGLTKTMPVLRYLMTKAGIDPPVVVETPK------------ATMTTTFESVRPGSEEQNL 107
YP_009337259      SPPAPPTFYEQLLEMG-------LERKKTIAVALFGAAAVGFWACKNRRRRVLLKPLS-------------------QESLVAGSAEMPA 103

YP_009337868      LAPNSQCRILLKNSDGDWVLTGCAVRVDLPIG-PFLVFPQHLCAG-----ELAAMSSRDEVIPLSLDGLEP---IAADLSMLPCP-SKLT 181
LSNVS1_PCR_ORF1   LAPNSQCRILLKNSDGDWVLTGCAVRVDLPIG-PFLVFPQHLCAG-----ELAAMSSRDEVIPLSLDGLEP---IAADLSMLPCP-SKLT 181
ASM94003          IHCRSQVKVLTR-DNTDWVIQGSAIRVATPDSKSYLIMPQHVCAASD--HVAVSDWKQKHVIEIDVDQLIP---IATDVCALPMTTLMET 223
YP_009330079      TAPRYQVIVCEK-KDGRLVKLGCAVRFDGN----FLVGPDHVLGEAD--LPKYAVGRQKF-VCLDKKERIP---LDTDLVAIRMSPSELS 154
YP_009329959      TPPKGQALVGFS-DGSSFMVVGCAVRMEDW-----LVMPDHVKSALGDRRLEIRSMDHKRVVTLHNAEVEAMQLVDTDLLAVQLSPARFS 187
AMO03214          TGPKCQAKIGYY-SDGQFVVVGCSVRFDDG----WLVGPDHVLG---G-NNKFAYGSQKRPVSLVGKEIVP---LATDLVAIKLTETEFS 125
QED21501          VHPPCQVFVGGM-VDGKFEVNGAGVRMYDY-----LVTPQHVVAPYSP-KVKIR-GLNGNDVEVDLLSFEG---LDTDICMIRLTPAQWS 186
AWA82253          TVPSFQCKIGTY-VSGQFSVLGSAVRFSTQSDKHYLVGPDHVLGESEG-EVKYAKGKQSE-ISLRGRERIP---LETDLVMIELTDKEMS 243
AMO03212          TCPKSQAVVCEK-VDGQFVMIGCAVRFENN----WVVGPDHVLGIGGG-RSKYLVGKQSN-ICLDLKERVP---LDADLVAIHLSDRDMS 151
QED21515          PFPKNQLLVGEL-VMGEFHAYGAAVRISDW-----LVLPAHVYSTTS--TPMVR-GKQG-IFDLTGYDFEE---IDTDLLAIKLDQKGWA 184
YP_009337259      LEPACQVAIAFESPDGTLVKVGSGVRGLVGCN-HVLFTAAHNLSFEQP----LWLIKGSKQVRLGDYSPIP---LAADAALLLVPEKTFG 185

YP_009337868      STGLASPTIGV-VRPGQRCVVSVVG-VDGLGTTGPLQ---TSP-VFGRLIYGATTNTGYSGAAYAS-ARTVYGVHTNGGKLNGGFALSYV 264
LSNVS1_PCR_ORF1   STGLASPTIGV-VRPGQRCVVSVVG-VDGLGTTGPLQ---TSP-VFGRLIYGATTNTGYSGAAYAS-ARTVYGVHTNGGKLNGGFALSYV 264
ASM94003          QLGARVASIGH-LPAGQTLTVQVVG-VGDRGTMAPIS---NGP-CFGSLSYTGTTLPGYSGAPYVMNGTTVLGIHTHGGKTNGGFALGYL 307
YP_009330079      TIGVSVAKIG--LVSAKGVYSQIVG-VDSKGTTGVLK---MDRVAFGRTVYDGTTLPGYSGAAYTS-GAFCSAIHQSGGAVNGGYASSYI 237
YP_009329959      EIGLAKVTVGPNLAEKFGAFSAIVG-AFGKGTTGTVK---HAQ-LFGKITYTGSTFKGYSGAAYMA-GTQLVGIHLHGGTVNAGYSASYV 271
AMO03214          SIGISVCKIA--PVPMQGIYSQIVG-PEGKGTTGVMR---NDIRSFGRVVYEGTTVGGYSGAAYTY-GSNAVGIHQMGGAVNGGYSANYV 208
QED21501          LLGTSKASISH-EIPAQGSYVSITGSESNLGTTGALR---EDPSVFGRVMYTGTTKKGYSGAAYAA-GKTLIGIHTNGGAVNGGYSASYI 271
AWA82253          TIGIQKAKIT--GVPSMGIIAQVVG-AMSKGTFGRLV---NDPMAFGRVVYEGTTVGGYSGGAYTC-GTAMLGIHQMGGNINGGLSASYV 326
AMO03212          TVGVSICKIG--PVPDCGSFVQIVG-PENKGTSGRLV---ADRSVFGRVVYEATTLPGYSGAAYTN-GPFIVAIHQTGGTVNGGYSASYI 234
QED21515          TIGTPVGTICH-EIGQYGSFVSIVG-ISGKGTTGVLQ---DDPHVFGRIIYHGTTKPGYSGAAYSS-GREIVGIHTNGGAVNGGYSASYI 268
YP_009337259      LLGVKVANFSP--LPGKSVAVSVTG-LAGKGTTGILSPVVNPHHGIGHVTYAATTAGGYSGAPYVS-GSQVYGIHIHGGMRNGGFEILYL 271

YP_009337868      YAMMKV-----------LKGVRDESSENWIHSVMADKSTRFLEMDYERVADSIVMRGDDGFYHAISGPAAEGLVKKLDKYKYDAG-MVDF 342
LSNVS1_PCR_ORF1   YAMMKV-----------LKGVRDESSENWIHSVMADKSTRFLEMDYERVADSIVMRGDDGFYHAISGPAAEGLVKKLDKYKYDAG-MVDF 342
ASM94003          HAMLKS-----------ATKVRDESSEDWARSLMS-GKDEGYTLTQVGEDTVVGTRD-GRYHVTS------------------------- 359
YP_009330079      WMLLKDHLRVEDVVEPESKKNWNSDTPEWLLSQFKAGKKLKWKRTG-DPDLIELMLDDGKFSRVSPASMHKAFGPTWHDHD-------VI 319
YP_009329959      LAMLKH-----------MFRIKDEGSDEWLEGMRRQGAEVVVDQDWRDMDECRIRVA-GRYHIIGVDTMSRVYGADWRRSG--------- 340
AMO03214          WILLKSQKIP-------ESKNWDSDTAEWISSQYKQGKKLKWKQGP-NPDEVEVFLD-GVYSMVQKSSMTRAFGNKWELSN-------EL 282
QED21501          ASLIQYE----------LRQHKVESSSDYLQTAMRNGMKIVVDGSWGHIDDVRVKVG-GRYHIMDKDSLKKSFGDMWELRVNEIS----- 345
AWA82253          WMLIKR-----------HAGVTEEDSEDWLLGQYKSGRRIRWKNSH-DPEYVEVETG-GKYSQVLKISMHKAFGQDWETSD-------FI 396
AMO03212          WMLLKDLSMTGETIIPEGRGNHNSDSSEWLLSQYNAGKKMKIRRTG-DPELAELYAD-GLYSRVTVDSMEKAFGPSWADKG-------FI 315
QED21515          LSMLNY-----------LDKKINEDSPGYLQNAFRKGKKIRVDKRWGGLDEIRAQVN-GRFAIFQRSSMRDAFGEDWEDMFDDVGRGSYK 346
YP_009337259      YALAKIA----------LEYTEEDTEDAFMERLFNSDDPDYYVQEVGDKVIVRVHGN--SHYHVVDAKKFRDYEARWENADYS------- 342

YP_009337868      RSKTGHTWSDIDPNHWESANPETAQVLASADTVPQGFHLDPSSRESPYIEELASQIRMFEDRLKSSLVDTALKKQIMDTFQQTLEPSRSL 432
LSNVS1_PCR_ORF1   RSKTGHTWSDIDPNHWESANPETAQVLASADTVPQGFHLDPSSRESPYIEELASQIRMFEDRLKSSLVDTALKKQIMDTFQQTLEPSRSL 432
ASM94003          -----------------------------------GDVAQQLKDRALGWKKLFSEVKAPKGSWADEEPE--------------------- 393
YP_009330079      EKGFDRSYRDVPRESILESVPCTSGSGEDHGSKLPGALSLLEPDQAWARLSLQQEMREFFNLSKRQQEDIRKCSQRQMSQNLASSGQAKT 409
YP_009329959      ----------------KRNLKTDRFVDLESCVLPGESSETLSGGSRVSVPSNPSLAELANQLTLHELEKLRERLALRAKDLRASATGSRA 414
AMO03214          EQAATRTYNDVARESILESVNP--VSGEVQSSKYLGVLNKLANAQGSEERSLLFETPEFVNLSKQQQKNIRSYLQLREQLMSTSSGLGSS 370
QED21501          -EKIKNGYSPESFVAKCPAPVSILESSGEATSLKSGASGLLEKSPEVEDQEKSTLIKLMQNMSKTQLKDSYLYILQREKTLNSQANVKEV 434
AWA82253          ERTANRTYDDVY-----ECVAT--PAGEANSSKRPGVSSVLHESRESAEQKPLRVIPGLQQLSKKQLADTRKYVQELLSQRQTSSGQAKS 479
AMO03212          QKATRATYRDVPNDILLESV-----SGEANGSSCPGASSILDNAQGSAKPSPQAETLESQKLSVKQRKAIAKSVMLAVQQSLISSGQEKT 400
QED21515          DGARKIGESKVPLSLPAIAEPAKVAAGEECTILNSGVLGSLTPSPDQQQSEKQALMRLLEPLSAEQLKDLYRRHLEFLRSSNSQVKSKSI 436
YP_009337259      SDDYVEECASEALNFQSPGQRSRAGPSIECLHRQPQLVTSSCQTSALPESSPSSDRRPQDPSNKRLKKLINSRLRRLQLKTNGTRSTPSR 432

YP_009337868      TLSLCRPELQHRLAKVMFGLH 453
LSNVS1_PCR_ORF1   TLSLCRPELQHRLAKVMFGLH 453
ASM94003          --------------------- 393
YP_009330079      MATKNS--------------- 415
YP_009329959      --------------------- 414
AMO03214          TSKTN---------------- 375
QED21501          KEKKNSGVLESTPSTSE---- 451
AWA82253          VLGTTSS-------------- 486
AMO03212          TDTTSGQVC------------ 409
QED21515          VVEKN---------------- 441
YP_009337259      SLVTQV--------------- 438
```

**Figure S4.** Multiple sequence alignment of deduced amino acids corresponding to predicted ORF2 of LSNV genome segment 1. Red-labelled letters indicate more than 60% amino acids identity.

```
YP_009337869     ------------------------------------------------------------------------------------------ 1
LSNVS1_PCR_ORF2  ------------------------------------------------------------------------------------------ 1
ASM94004         ---------------------LLPSFSGDGGPFPPPPDPPPPPGFIRDGVAQMGGQFVRRRAAVDPEIDGCLQALFPDELYSAAVHGYVP 69
ACJ12846         ------------------------------------------------------------------------------------------ 1
YP_009336757     ------------------------------------------------------------------------------------------ 1
YP_009342317     ------------------------------------------------------------------------------------------ 1
QHA33876         ------------------------------------------------------------------------------------------ 1
ASA47392         ------------------------------------------------------------------------------------------ 1
QHA33888         ----------------------------------------------------------------------------------------ME 2
YP_009330080     ------------------------------------------------------------------------------------------ 1
APG75843         ------------------------------------------------------------------------------------------ 1

YP_009337869     ------------------------------------------MEEAYAAVRWDIPSDFLSREHYERTLLRLDLQASPGYPYLREAATIGL 48
LSNVS1_PCR_ORF2  ------------------------------------------MEEAYAAVRWDIPSDFLSREHYERTLLRLDLQASPGYPYLREAATIGL 48
ASM94004         ADPTTGAVSRSFETQAKIAAGARRSYDRPTPVELDTVL--AWTESKYSALRHALPDDFLSKDHFVRCLHRLDLTSSPGYPFMREASTNGQ 157
ACJ12846         ------------------------------------------------------------------------------------------ 1
YP_009336757     --------MKSFAVQSRTAHIVRERTISPTDEELEHILYHAEQ--ALAPAKWKLPSNWNSRTAFEAALAELDWQSSPGYPLLREAPTIGD 80
YP_009342317     ---------------------------------------------------------------------------------MREAPTIGK 9
QHA33876         ---------------MTNAAALRAALVEPSEMEEEAVLYYLERKYKTAMDKRPVPKDWACRSRFEILLLYLDNASSPGYPYMREKPTIGE 75
ASA47392         ----------------------------------------------MQHPKGRVPENWNSRERFMEILKDLDFSSSPGYPYMREAPTIGR 44
QHA33888         TYCLKAVEEPDQELKDRILY---------------------IMEKAYSTVIWTLPDNYDSYLAFERAVSNLDLTSSPGYPYCLEKPTIGQ 71
YP_009330080     ------------------------------------------MEQAYSPVVWSLPDDFDSRTRFDWAIRRLDMQSSPGMPYMREAPTNGK 48
APG75843         -----------------------------------------------MHSGYVIPADWNSKSRFMTLLLTLDYSSSPGYPYLREAPTIGK 43

YP_009337869     WLGVTLEGQFAPDQVERLWQHILLILSGEFDHYYRVFVKDEVHRKKKADEGRWRLILASALPMQVLWHMLFAPMNDLEAEKIFHTPSAFG 138
LSNVS1_PCR_ORF2  WLGVTLEGQFAPDQVERLWQHILLILSGEFDHYYRVFVKDEVHRKKKADEGRWRLILASALPMQVLWHMLFAPMNDLEAEKIFHTPSAFG 138
ASM94004         WLGWTFQ--PDPVRVDLLWEYVKQVIRGKFEHLYRVFVKDEVHKTSKAEAGRWRLILASALPVQMVWHMLFGDMNDMEAAKVFETPSLYG 245
ACJ12846         ------------------------------------------------------------------------------------------ 1
YP_009336757     WLLIKGTLERDPSKVERLWDMVQKVLKGEYIHYWRVFIKSEPHKRSKALEGRWRLITAASLPVTVAWMMTFKKLNDRLSHYDAALPVQQG 170
YP_009342317     WLK-WELDHPDPIQACRLWIMVQQVFAGTYDHYFRVFVKDEPHKIAKAREGRWRLIIASSLPVQVAWNMAFRAMNDLLIKKVYRTPALQG 98
QHA33876         WLGADGFGGFNTVQVERLWYDVQQVLSGNYEHLFRVFVKDEAHKKAKADTNRWRLIVASSLPVQMVWRMLFHEQNLALNECSSEIPSKHG 165
ASA47392         WLKTDEVSKFDPVQVERLWYDTQKVMAGTYDHYFRAFVKDEPHKKAKAEQSRWRLIIATSLPVQMVWRMLYREQNDAMNKLHKHLPSKHG 134
QHA33888         WLEHDGL-HPSWRAMQRLWIDVQRVFNGTFDHIQKVFIKMEPHKVSKIVEQRWRLIIASALPVQVVWHMLFDYQNDKEIEQAYYIPSQQG 160
YP_009330080     WLKWDGV-EYDAMQANRLWHDVQCVLSDDWEHVIRVFIKQEPHKKHKAREGRWRLIMASSLPVQLVWHMLFSYMNDLEISECYNIPSQHG 137
APG75843         WLRTDECGSFDPVQVERLWYDVNMVMAGSYEHLFRAFVKDEPHKIAKAKENRWRLIIASSLPVQMVWRMLYTGQNEALNKYHDWCPSKHG 133

YP_009337869     VSFVYGEWKMFKNYCRVKRLEVAIDKSGWDWNAPGWVFDADLQLRYRLCNQAERPAGHLWFALARKLYDDAFEHSRCLLPSGQVYVQEFS 228
LSNVS1_PCR_ORF2  VSFVYGEWKMFKNYCKVKRLEVAIDKSGWDWNAPGWVFDADLQLRYRLCNQAERPAGHLWFALARKLYDDAFEHSRCLLPSGQVYVQEFS 228
ASM94004         MSFCAGGWKYFKDYAASKGLTHAVDKSGWDWNAPGWVFQADLEVRRRLCVNVT----DEWCALADAMYRDAFLDSRLVMPDGTIYKQQFA 331
ACJ12846         ---------------------------------------------------------------------------RCLLPSGQVYVQEFS 15
YP_009336757     YVWCSGGWKNFKRRIEQDGLTISVDKKAWDWGAPGWAFEADLELRTRLCLNPS----PEWHRVATLLYADAFQTAKLLLPNGIVYEQQFH 256
YP_009342317     LILCYGGWKRFLAYAQTNNWTMSKDMSGWDVNAPGWVFDADLELRTRLCSNPS----ASWIRLVRMLYDDAFRDAKLLFSNGLVYQQQFS 184
QHA33876         YVHCYGGWRQFLAEAKTAGMKYSRDISGWDVGAPGYIFRIVGKMRQRWKG-VT----ASWITVQERMYRDAYEESNLIFSNGIVVKQEFS 250
ASA47392         FVFCYGGWMDFVAEAKSKGLKVSRDISGWDVGAPGWVFKVVGALRENWGG-VT----ASWIRVHRLMYKDAYSEAKILFSNGIVVQQEYD 219
QHA33888         LKLCGGHWKLYYQQWKQRGFDVGLDKSAWDWTAPGWALMLDLKFRLRMGRGSR---MREWATTAAALYRDMFETSKLLMPEGYLLQQEFC 247
YP_009330080     LILVGGGWKDYLRSWKEEGLSVGLDKSAWDWTAPRWVMDWDLDFRYRMGRGKR---MEEWHRLAKLMYHHMFDHPVLQLSDGTLLRQTVP 224
APG75843         FVFCYGGWMKFIAQAKTKGLNVSRDISGWDVGAPGWVFEVVGAWRESWPG-AT----DSWIRVHRMMYDDAYKNSRIIFSNGVVVRQLFG 218

YP_009337869     GFMKSGIVNTISTNSHAQIMLHMLACKRSGEPVTP-------ILACGDDTIQAATSAGYIEELAKAGCIVKSVDRKLEFMGFNFE-GAMQ 310
LSNVS1_PCR_ORF2  GFMKSGIVNTISTNSHAQIMLHMLACKRSGEPVTP-------ILACGDDTIQAATSAGYIEELARAGCIVKSVDRKLEFMGFNFE-GAMQ 310
ASM94004         GFMKSGVVNTISTNSRAQWMLHCLASLRVGTQPSP-------LVACGDDTLQSQYPPGYLYELGRAGCLIKHCKESLEFMGFSFD-GKIV 413
ACJ12846         GFMKSGIVNTISTNSHAQIMLHMLACKRSGEPVTP-------ILACGDDTIQAATSAGYIEELAKAGCIVKSVDRKLEFMGFNFE-GAMQ 97
YP_009336757     GFMKSGCFNTISTNSNCQILLHYLAEYRWAKAEGVSIKEHSKILACGDDTLQAFYTPLYGELLEQAGCTVKEAAQNTDFMGYNYD-GPPE 345
YP_009342317     GFMKSGVYNTISTNSNCMFFLHVIASRRARIPLRP-------IGAVGDDTIQSPFPDAYIDHLQSLGCVVKEQNVGLEFIGTDFRSGEPR 267
QHA33876         GFMKSGLFNTISDNSISMVGIHALACLRASIPIGS-------IFATGDDVLQSTISDNYLDELGKLGCRVKEVLYHIEFMGVNYSSGKPE 333
ASA47392         GFMKSGLYNTITDNSLSMGAIHGLACKRSGLPFGR-------YVVTGDDIVQSHVNEAYLEELYTLGCRVKEVLHHLEFMGINFSSGKPE 302
QHA33888         GLMKSGCVNTISTNSHMQVMIHIAANQHGKV--YPLPS------CCGDDTLQCSTQCDDLEKYRRYGIFVKQVERGLEFVGHRFEHDGPK 329
YP_009330080     GIMKSGCVNTISTNGHAQVMMHCVVAEDSNVPYEPYPK------TCGDDTLEHPMHTQSLEYYGRYGVVVKSVSENMEFVGHEFTDSGPH 308
APG75843         GFMKSGLFNTISDNSLAMGAIHSLACVRSGLPFGH-------YIVTGDDIAQSTVSAGYLDALYKLGCRVKEVLYHLEFMGVNFSSGKPE 301

YP_009337869     PIYTVKHIASFTYKSEELHGEILDSMCRMYA-HHPWFDCWKMLAGSLGHNM-KSRAWYQFFLDSTANIRITKSI- 382
LSNVS1_PCR_ORF2  PIYTMKHIASFTYKSEELHGEILDSMCRMYA-HHPWFDCWKMLAGSLGHNM-KSRAWYQFFLDSTANIRITKSI- 382
ASM94004         PLYEEKHVATVRYKAQVIS-EVLDAYCRMYA-HAPLYDSWAKLARKLGVSL-MSRSFYRTFMDSPYPLRV----- 480
ACJ12846         PIYTMKHIASFTYKSEELHGEILDSMCRMYA-HHPWFDCWKMLAGSLGHNM-KSRAWYQFFLDSTANIRITKSI- 169
YP_009336757     PMYFAKHLVKFALQKPALRPETLDSYMRMYA-TSESRHFWRAVANELEIHL-MSDSYYARWANVPEP-------- 410
YP_009342317     PMYFDKHLVSICN-TEDVP-STLDSFCRLYAHEPDKLAFWLELGRQLGVSL-RSSAYYQFWYDSPWARMLEW--- 336
QHA33876         PMYIQKHLFNVALKTDVLE-ELLDSYCRLYS-ESKNFPFWAKLAEDLGINV-RSQRYYQFWYSSPMGAMVAKWLS 405
ASA47392         PMYTQKHLYNMTVKSEYVA-EVLDAYCRYYA-ESRKYGLFEEVAKQLGVPV-RSRWYYQFWYSSPLAAIYYGLA- 373
QHA33888         PAYYSKHLYKACYQEDDILPQYLDSMCRMYV-KTPQFEVWENIASELGIKL-FSREYYMYWYDVATD-------- 394
YP_009330080     PLYISKHMKKLQYLADDIIPDFLDAMARMYV-HTRYFSIWEDLAIVNGTPLPMSKMAYRYWYDFSV--------- 373
APG75843         PMYTQKHLFNLLTKEQFIA-ETMDSYCRYYA-ESSKYHWFTEIAKELAVPV-KSRWFYQFWYSSPLAEIYHKLG- 372
```

**Figure S5.** Multiple sequence alignment of deduced amino acids corresponding to predicted ORF1 of LSNV genome segment 2. Red-labelled letters indicate more than 60% amino acids identity.

```
YP_009337863      ----------------MTNQHVCPICKRSFRTAQGLADHKRDRHEACSTGPAPKPQRVRPKRE----QRQAAPVSVPTSSGPARYSNRVT 70
LSNVS2_PCR_ORF1   ----------------MTNQHVCPICKRSFRTAQGLADHKRDRHEACSTGPAPQPKRVRPRRE----QRQAAPVSVPTSSGPARYSNRVT 70
ASM94005          ----CGSFLDQTDPGLTAMVFKCPKCSRQFKTKAGLSQHEVDKHTSAP------PQSVR--------RRQSAPVSLATTAGPTRAAGTVR 72
YP_009333381      ---------------------MAKKPTKKNARNRGSNRP----------------------------RGAPVAQATNIHVGSRDPP---- 37
YP_009337277      ---------MVG-------SVECEYCKKKFKSKRAKNQHVSMVHKINEKMAPPAVAKTGRTRGSLRLRRTRTSNLPGFDIAPSRVP---- 70
YP_009345111      -------------MNAQTKKITCHACGRRFNTQQALHQHQQAVHGAANQRVPRGAGRGGRGRR----NGQNNQGPARQGTAQSVTSR--- 70
AWA82259          RSVLPNFLPLFFKMSESAKKFVCPVDQRRFLTKAALTQHEIASHGKAHNSPKSSGHPNSKPRRP---VQQGAPVSVPSVVRGGGSD---- 83
AXQ04819          ---------MPSKKKNVN-SCVCPHDGKQFSTRAALQQHKKMVHASPNETRPAPQLRR-RPR-----RRSGNGGGLSSDLAPSRVP---- 70
YP_009182197      ---------MPSRKKNVN-SCVCPHDGKQFSTRAALQQHKKMVHASPNETRPAPQPRR-RPR-----RRSGNGGGLSSDLAPSRVP---- 70
QIJ25876          ---------MPNNKRNNNNGCICPHDGKHFATRAALLQHKRMVHQGVSETCPSTQPRR-RNR-----RGPRNGGAITSDLAPSRVP---- 71

YP_009337863      LQGSNERFRTDDYP---ASRLKTG-TLLTTIPVSPDLVPRLSTQAKAYQRIKYHSVALHIDAKGSTAQSGGYLAVFISDITDEEISLERA 156
LSNVS2_PCR_ORF1   LQGSNERFRTDDYP---ASRLKTG-TLLTTIPVSPDLVPRLSTQAKAYQRIKYHSVALHIDAKGSTAQSGGYLAVFISDITDEEISLERA 156
ASM94005          LWGRDERLFTADYDGQGPTAMKTG-QLIQRIPLVPGIVQRLETMARGFQRIKYDKVSLEIDAKGSTSVSGGYLAVFVSDPSDEPD-LARA 160
YP_009333381      --IVVRGTGRFAHSTHAANSLSDG-DVLFDSIINPSSFPRMSRIASAYQRYRFTRLAFTIQPMCPATTGAGYVAGFLKDPTDEDTSFDAI 124
YP_009337277      -TVRGGMINISGEDRIGAFDLKSGKAVFMSVDISPSMSARLTTIARAYQRIKWNAVKIVVTPQASAMTNGGYVCGFIADPSDRAVTASDL 159
YP_009345111      -ATPNQGVSVSGFERLGSIDIVNA--ASASWQVNAWMTPRLQAMSSAFQRIHYLSLKVNVVAFGSAIGTGGYVIGFVADPSDATPTISQL 157
AWA82259          --LARLSGVDRVYHGEIATNMDQG-KPVFSLLITPGTFSRLKTVSLAYQRIRYLSVNFRIETQISTTTSGGYVVGFVRDPADVVDDINTL 170
AXQ04819          -TVRNSAITTSGSDKVLVFTIKAQQSLIKSIPIHPGSIPRLSNLAKSFQRITWRSLAVHVSPQVSTSINGGYVAGIVTDPDDEHITASDL 159
YP_009182197      -TVRNSAITTSGSDKVLVFTIKAQQSLIKSIPIHPGSIPRLSNLAKSFQRITWRSLAVHVSPQVSTSINGGYVAGIVTDPDDEHITASDL 159
QIJ25876          -TVQSSSLTTSGSDKVLVFTIKAQQSVIQSIPIHPGSIPRLSNLCKSFQRITWRSLVVYVSPQVSTTINGGYVAGIVTDPDDEHITSTDL 160

YP_009337863      GSFGGSVGKKYWENATVTATNLTPLFYTSRGEDERLWSPGYFAIYCDGSSNTAISLTYRLTWSVTLSVPSSE-LKQADVITT--MVDLWP 243
LSNVS2_PCR_ORF1   GSFGGSVGKKYWENATVTATNLTPLFYTSRGEDERLWSPGYFAIYCDGSSNTAISLTYRLTWSVTLSVPSSE-LKQADVITT--MVDLWP 243
ASM94005          GAYSGAVGRKWWENSTISATNLSPLFYTSLSREVRTASPGYFAVFCDGKPSQDVSLTYRLTWETTLSVPSMEDEATLREVDV--VVDLWS 248
YP_009333381      QGSQGAVIGKWWEHKRIEVRPPTDLLWTSLGENPRLFSPGKFVMTAVGTNTDAVNVSVLCEWEAVLSVPSLEDYSEKPVTEYFSTKDLLN 214
YP_009337277      SASQGAQTKKFYETAVVYMPRKTDLLYTSAGEDPRLFMPASFWIISEGLPSSNLTMIVSVVWDVTLSQPTLENSHNNSFLLVGEIVPNPA 249
YP_009345111      QSQAGARTCKSWENCDVPLVVPGTLYYTSRDNEPRWYSPGRIVILVDGIASAGGKLVINLHWSVRLSVPTATPPTVLKDIFLSEHLYVIP 247
AWA82259          MSQKGSVATKWWQSSTVAATPTNRLYYTSESVELREFSPGRLVMMVDGAATQRGSFTVYANWTAELSVAGLESNAQVKREIVAISNWYTR 260
AXQ04819          SSTAGSQTKKWYQDAIIRMPLKTDKLYTSAGEEPRFSSPGTFWICSEGNPSSDLTVVVTFRWTVHLEGSTLEDDATSSFVLSGSLNSIGG 249
YP_009182197      SSTAGSQTKKWYQDAIIRMPLKTDKLYTSAGEEPRFSSPGTFWICSEGNPSSDLTVVVTFRWTVHLEGSTLEDDATSSFVLSGSLNSIGG 249
QIJ25876          SATAGSQTKKWYQDAIVRMPPKSDKLYTSAGEEPRFSSPGTFWICSEGTPSSDLTIVVTFKWKVHLEGSTLEDDTSSSFVLSGSLNSIGG 250

YP_009337863      TAGKASFSDKDGSEE---NLFEPSLPVGTVVRFPYAVG---IEYKEGAGDTGTVPIMFAVVTASSG--LKASNDRVDTSVVWQSDVSR-R 324
LSNVS2_PCR_ORF1   TAGKASFSDKDGSEE---NLFEPSLPVGTVVRFPYAVG---IEYKEGAGDTGTVPIMFAVVTASSG--LKASNDRVDTSVVWQSDVSR-R 324
ASM94005          KAGSAAVATEGGVQKPLVDLISPVPPVGTVLRLPFPVS---VEYSEGSGDTGTIQAMFLIVTSDGK--LTLSNDRSTAVKQWQSDLAI-R 332
YP_009333381      ----SSYVHNSNSILQTNVFDGDVIPTGTILRIPFPID---VDYSLGAGDVASARYCYFLAGSHNLTWGLYSGGSFSGGSNYYSDGQPTQ 297
YP_009337277      NYNLRYSPPGG-DPQDDASSIIPAVLRETPGYHYFRVPTFTIEYSEGTGDTGTIQAHFVVYHTTDKKLYYSSNGRDVETTMWQGNVDAHQ 338
YP_009345111      GKNILGWKDGTTFRG----NISKFLSTPVPGDSWLSVPTFGIEYNEGVGDTGTILIHYLKYNSSEGSLTCSNDGEKVVTQVWQASVAA-Q 332
AWA82259          SGHQGFWTNQTDRAS---SRLIDLIGADAITGRTYRLP-HSVTITQENEAVRNCNWVYAQDDNTLLPAYASPTDKDTGALKINDLILVKG 346
AXQ04819          KSIMGYKACGSTTVEEDVSGIIPAPIREVPGTHFFRVPTYTVEYQEGVGDTGTIQMHFVAYKTDTKNFHFSEDGSSVNASDWQADVDN-Q 338
YP_009182197      KSIMGYKACGSTTVEEDVSGLIPAPIREVPGTHFFRVPTYTVEYQEGVGDTGTIQMHFVAYKTDTKNFHFSEDGSSVNASDWQADVDN-Q 338
QIJ25876          KAIMGYKACGSITVDEDVSKLIPASIKDVPGTHFFRVPTYTVEYSEGVGDTGTIQMHFVAYKTDTKNFHFSEDGSTINASVWQNTVDN-Q 339

YP_009337863      LLYPKSIRLIVEKRPGEVSRAGQVSQFSPCPPTSLASSETPSTETVGNNENRPTCSVRFTTPSSSDLMKSMHGCKTSMSDLRARPVCTVS 414
LSNVS2_PCR_ORF1   LLYPKSIRLIVEKRPGEVSRAGQVSQFSPCPPTSLASSETPFTETVGNNENRPTCSVRFTTPSSSDLMKSTHGCKTSMSDLRARPVCTVS 414
ASM94005          LLIPHSVSLIVDQ--GEDPRSGLATHFSRSLRSSGKSGTMEQHSSTKCLEISGACSK-------TDLSRSSQ-NWTAKHQHVAR-----S 407
YP_009333381      CVLPKGTRFEVIPN---------------------------------------------------------------------------- 311
YP_009337277      TLVPCGTFMKYVGQENSCRDTKLLTPKLLSETSPGSTSSSTNLSAKMQ------------------------------------------ 386
YP_009345111      VVVPRNTILTIVDYNKPAKPAVSDFIQPLVSNFNNVGIDSSRNSTTPS------------------------------------------ 380
AWA82259          SVLVDVTPTTVKSGEEVAPSSSVCQTMTAESLMPLLEFVR-------------------------------------------------- 386
AXQ04819          VLCSDGTYFKYTGQGNACKVASTVAPR--SSPSHLKRHSREKLMSLSR------------------------------------------ 384
YP_009182197      VLCSDGTYFKYTGQGNACKVASTVAPR--SSPSHLKRHSREKLMSLSR------------------------------------------ 384
QIJ25876          VLCSDGTYFKYTGQGNACKVASTAVLR--SSPSPLKRHSRDKLQTLSR------------------------------------------ 385

YP_009337863      STLPPSPNPSQESILLTFLESQVPRYQAARIVSQEVKTHEFPLLHHRNHLRLEMHWDLVPLMRGRRIPVYELSRHFDDDFSLKFTINNTV 504
LSNVS2_PCR_ORF1   STLPPSPNPSQESTLLTFLESQVPRYQAARIVSQEVKTHEFPLLHHRNHLRLEMHWDLVPLMRGRRIPVYELSRHFDDDFSLKFTINYTV 504
ASM94005          SLLEPSPK--EASIRLSPSPLSEARLACATQMAGAGRIFAKGLCVEGKSWELTVSPSIDGLM--------EHPPEWYNEFWLR-TARGSV 486
YP_009333381      ------------------------------------------------------------------------------------------ 311
YP_009337277      ------------KLETLFLELKQNYQKDSKPLMEESMILKPNLVEKLEQLEL-------------------------------------- 426
YP_009345111      -----------AGETGTDPPYSTYLELLRRQSALDSKISRIELQLSKMVKVLMPKDQ---------------------------DSSQSS 432
AWA82259          ----------------LLESFRVTSETLPKSSASSSRQSQCSRDLSDSTEKS-------------------------------------- 422
AXQ04819          ------------KVASLEASLKK-FEMFSQ---RDSMTSSPELIPSPENLEL-------------------------------------- 420
YP_009182197      ------------KVASLEASLKK-FEMFSQ---RDSMTSSPELIPSPENLEL-------------------------------------- 420
QIJ25876          ------------KLASLEASLKK-FEMLSP---RDSMTSSLELIPNQVSLEQ-------------------------------------- 421

YP_009337863      SFREVVEAVEVLNRGRYFCLGFIALCLQHHFGADSSEVVPEEFPSVWYDESFHYRLPCRSCHDNMESDYSFEELDPEVEYD 585
LSNVS2_PCR_ORF1   SFREVVEAVEVLNRGRYFCLGFIALCLQHHFGADSSEVVPEEFPSVWYDESFHYRLPCRSCHDNMESDYSFEELDPEVECD 585
ASM94005          TCNSPARAAEVLPTLRRLLLS----------EYDIDDVLDSSMPS-----------------SDPEDVGGFEVLG------ 534
YP_009333381      --------------------------------------------------------------------------------- 311
YP_009337277      --------------------------------------------------------------------------------- 426
YP_009345111      RSRSPSKDTTKLDSKGT---------------------------------------------------------------- 449
AWA82259          --------------------------------------------------------------------------------- 422
AXQ04819          --------------------------------------------------------------------------------- 420
YP_009182197      --------------------------------------------------------------------------------- 420
QIJ25876          --------------------------------------------------------------------------------- 421
```
